# Functional and structural basis of extreme non-coding conservation in vertebrate 5’UTRs

**DOI:** 10.1101/2020.06.29.165878

**Authors:** Gun Woo Byeon, Elif Sarinay Cenik, Lihua Jiang, Hua Tang, Rhiju Das, Maria Barna

## Abstract

The lack of knowledge about extreme conservation in genomes remains a major gap in our understanding of the evolution of gene regulation. While previous findings have mainly focused on the role of extreme conservation at the level of DNA in transcriptional regulation, its implications for RNA biology remains largely unexplored. Here, we reveal an unexpected role of extremely conserved 5’UTRs in translational regulation that is linked to the emergence of essential developmental features in vertebrate species. Endogenous deletion of conserved elements within these 5’UTRs decreased gene expression at the post-transcriptional level. A large-scale reporter library of extremely conserved 5’UTRs revealed the widespread presence of cis-regulatory elements that promote cell-type specific regulation of translation. As these elements function as RNA molecules, further understanding of their potential structures was essential. We therefore developed in-cell mutate-and-map (icM^2^), a novel methodology that maps RNA structure using high-throughput mutational analysis, previously impossible to perform inside cells. Using icM^2^, we determined that an extremely conserved 5’UTR encodes multiple alternative structures whose relative proportions are actively maintained by ATP-dependent RNA helicases. We further show that each single nucleotide within the extremely conserved element maintains the balance of alternative structures important to control the dynamic range of protein expression. These results explain how extreme sequence conservation can lead to RNA-level biological functions encoded in the untranslated regions of vertebrate genomes.

## Introduction

One of the most fascinating findings from the comparative analysis of vertebrate genomes is the existence of extreme sequence conservation in some non-coding regions, at levels often greater than even coding regions with perfectly invariant polypeptides^1–3^. Historically, the earliest whole genome sequencing efforts in humans and mice revealed “ultraconserved” elements, which were defined as any ≥200 nucleotides (nt) stretch with 100% sequence identity between the two species^4^. It has since been shown that these regions are undergoing strong purifying selection in humans and are not merely mutational coldspots^5^; thus their perfect sequence conservation led to the widespread conjecture that they are required for organismal viability. As the number of species with completed genomes increased, various finer criteria and terminology have been used to define such exceptionally conserved sequence elements^4, 6–12^. However, the acceleration in our ability to sequence and annotate more genomes has not been matched by our ability to ultimately explain the fundamental problem initially raised a decade ago: why does such extreme conservation arise during evolution, and what are the functional roles for such sequences in the genome?

To date, efforts to understand the phenomenon of extreme conservation have heavily focused on intergenic sequences. Transgenic reporters of highly conserved intergenic elements frequently display spatiotemporal specificity in their RNA expression patterns, suggesting possible roles as developmental transcriptional enhancers^13–15^. However, early *in vivo* knockout studies paradoxically yielded viable mice lacking grossly deleterious phenotypes, raising uncertainties about the relevance and contribution of highly conserved elements on organismal development^16, 17^. Only more recently have mice with loss of single or pairwise deletions of ultraconserved enhancer elements been shown to produce more subtle developmental phenotypes such as neurological defects or growth abnormalities due to their impact on the transcription of neighboring genes^18^. In combination with another finding that enhancer redundancy is a widespread feature of mammalian genome architecture, this work has pointed to the idea that finer defects may have profound consequences for evolutionary success in the wild over long timescales of hundreds of millions of years^19^.

However, beyond its significance in transcriptional regulation, the biological meaning of extreme conservation in post-transcriptional regulation – i.e. the level at which the sequence is expressed as RNA molecule – remains largely unknown. While few examples - such as the functional roles for ultraconserved regions transcribed as long non-coding RNAs or alternatively spliced poison cassette exons - have been described^20–24^, RNA-level mechanisms for extreme conservation have not been explored widely. In particular, the potential for extreme conservation to elucidate the functions of 5’UTRs in the translation of vertebrate mRNAs and in the context of organismal development and disease deserves special attention. Most of our current understanding of the role that 5’UTRs play in translation centers around the framework of the cap-dependent, scanning model of translation initiation^25^. The dominant parameters of translation efficiency in this model are the length, global structuredness, and competing start codons of the 5’UTRs, properties that can be retained across much sequence variation. The observation of extreme sequence conservation across extended stretches of 5’UTRs therefore suggests the possible presence of novel and more specialized translational cis-regulatory elements.

In a paradigmatic example, the Hoxa9 5’UTR contains a ∼650nt extremely conserved region that has previously been shown to mediate non-canonical translation initiation through a structured IRES-like RNA element^26^. Knockout of an ∼150bp functional element within this conserved region in mice results in diminished spatio-temporal Hoxa9 protein expression and a pronounced axial skeleton phenotype leading to a homeotic transformation, demonstrating how 5’UTR RNA sequences important for specialized translational regulation in the developing embryo can undergo extraordinary negative selection. We thus were inspired to ask if there could be a broader, systematic trend for extreme conservation to reveal currently unknown translational regulatory sequences, and conversely, if such regulatory sequences could help to explain the functional basis of extreme non-coding conservation in mRNAs.

To this end, we begin our study by defining a set of “hyperconserved 5’UTRs” (a term we use to refer to extreme conservation in 5’UTRs under the criteria described below) across vertebrate genomes, creating a catalog of sequences with exceptional conservation patterns for further systematic characterization of their functional and molecular features. We address whether the 5’UTR hyperconserved elements can impact translation of mRNA transcripts in cis by deleting them from endogenous loci as well as by dissecting them through reporter assays to facilitate mechanistic understanding. This led to the discovery of cis elements that contribute significantly to the translational control of multiple genes known to be important for embryonic development. We describe how such functional roles of 5’UTR hyperconserved elements may, at least in part, be reflective of the widespread occurrence of non-canonical translation initiation regions. We then ask what such exceptional conservation patterns may mean for the RNA secondary structures of the 5’UTR, using a novel technology we develop called in-cell mutate-and-map-seq to resolve the structural ensemble of a hyperconserved element in its native cellular context. This revealed the unexpectedly rich landscape of alternative structures previously inaccessible through purely computational or conventional experimental approaches that may explain the near perfect conservation of 5’UTR hyperconserved elements. Together, our data lend support for the critical importance and utility of extreme conservation in advancing our functional understanding of eukaryotic translation beyond canonical models and in discovering functional structures encoded by mRNAs of vertebrate genomes conserved across evolution.

## Results

### Defining hyperconserved 5’UTRs in vertebrate genomes

The function of extreme non-coding conservation has not been investigated for mRNA 5’UTRs, where such exceptional conservation may have resulted from critical post-transcriptional regulatory mechanisms under negative selection. We hypothesized that extreme conservation could reveal sequence-specific cis-regulatory elements important for translation that remain poorly characterized in vertebrates. To address this, we used the conservation pattern of the aforementioned Hoxa9 5’UTR as our archetype in selecting a set of other 5’UTRs in the genome. The length of the extremely conserved stretch in Hoxa9 5’UTR is ∼650nt; the size of the functional element within the conserved stretch is around 350nt^26^. We thus aimed to find 5’UTRs that had a highly dense run of matching nucleotides across the multiple species alignment (MSA), ranging in the order of several hundred nucleotides. We used PhastCons, a phylogenetic hidden Markov model based method designed to identify such blocks of conserved regions given a MSA^27^. PhastCons assigns each identified conserved element a log odds score (LOD) that reflects both conservation probabilities of nucleotides and the length of the element. Starting with PhastCons elements for a 60-way vertebrate MSA, we chose a LOD minimum of 500, which marked extremely conserved regions throughout the genome that are 100nt long, on average. Using mouse RefSeq gene annotations, we intersected mouse 5’UTRs with the LOD≥500 PhastCons elements (representing the top 8.25% of all PhastCons elements identified in the genome), requiring at least ≥250nt overlap. This resulted in a set of 589 5’UTRs for 499 genes (**Fig. 1A, Table S1**). Similar to the set of “highly conserved elements’’ used in the PhastCons paper, the elements here are longer but less perfectly identical than ultraconserved elements^4, 27^. The median nucleotide identity between mouse and human genomes in the conserved regions in the selected 5’UTRs is 92.3%. The average total length and the average number of nucleotides overlapping PhastCons elements for these 589 5’UTRs are 674nt and 389nt, respectively, and they tend to be found more frequently closer to the start codon than to the 5’ end (**Fig. S1A, S1B, S1C**). For the remainder of the text, we will refer to these 589 5’UTRs as hyperconserved 5’UTRs (h5UTR) and the LOD≥500 PhastCons elements within the h5UTRs as 5’UTR hyperconserved elements (HCE). h5UTRs serve to provide us with a testbed for assessing the functional and molecular features of extreme conservation occurring in the 5’UTRs.

**Figure 1.**
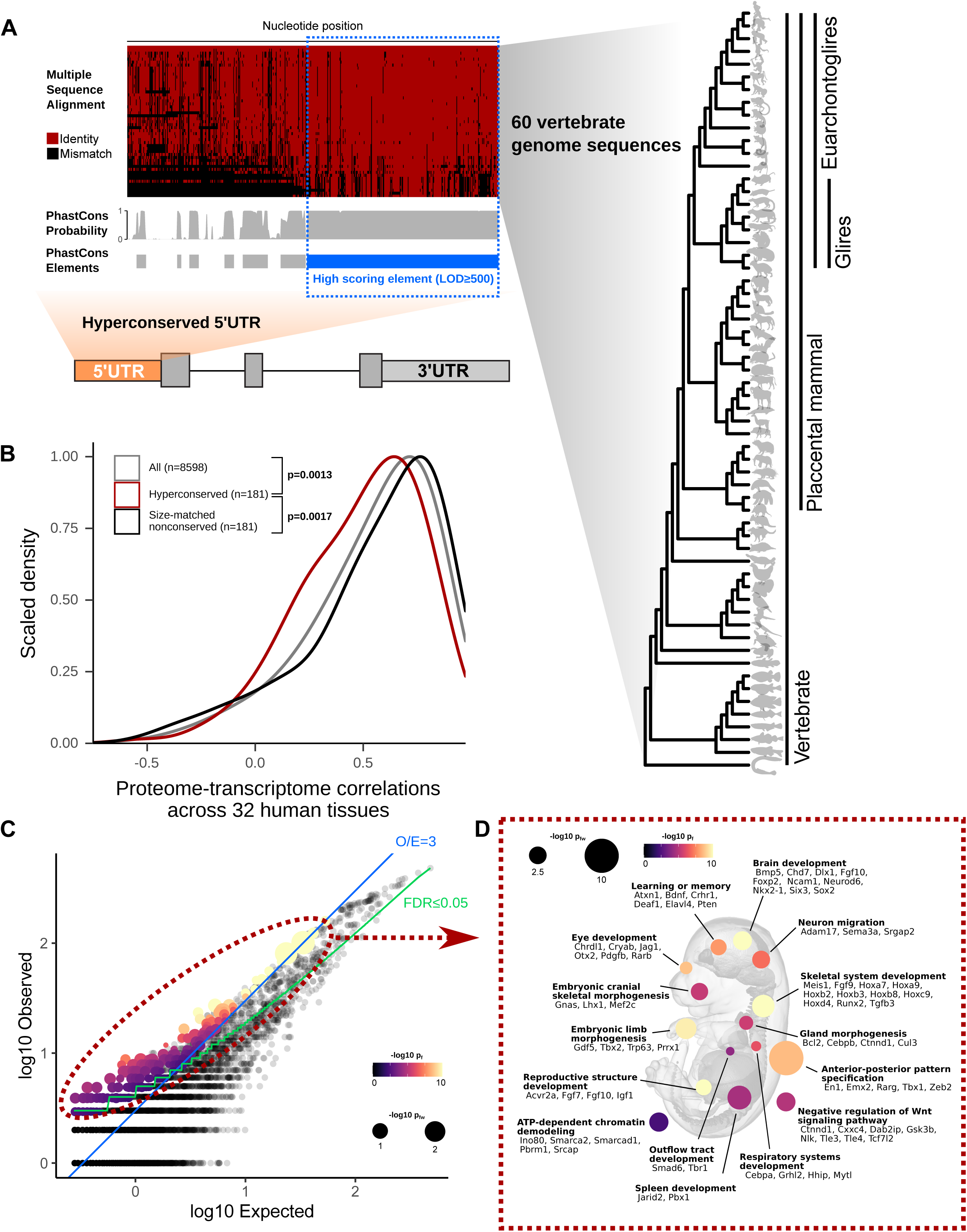
**Defining hyperconserved 5’UTRs in vertebrate genomes** A) Schematic illustrating selection of hyperconserved vertebrate 5’UTRs. We begin with 60-way multiple species alignment of vertebrate genomes, its per-nucleotide PhastCons probabilities, and conserved element prediction tracks. High scoring (≥LOD 500) PhastCons elements are overlapped with RefSeq annotated mouse 5’UTRs. We define those with overlap ≥250nt to be hyperconserved (also see **Table S1**). B) Distributions of cross-tissue transcriptome-proteome correlations (GTEX consortium data across 32 human tissues) for all genes, genes with h5UTRs, or genes with size-matched non-conserved 5’UTRs. Indicated p-values are from Wilcoxon rank sum tests for cross-tissue correlation values between h5UTR genes and all genes or between h5UTR genes and size-matched non-conserved controls. C) Scatter plot illustrating the term enrichment strategy and criteria. X-axis and y-axis plots expected and the observed number of genes for each term. Blue dashed line indicates the minimum observed/expected ratio cutoff of 3. Green line indicates expected and observed counts where Fisher’s test p-value (p_f_) is estimated to have FDR=0.05. Neighbor-weighted test p-value (p_fw_) ≤0.05 is further used as an additional cutoff. The final set of enriched terms passing filter is colored by p_f_ and sized by p_fw_. D) Visualization of representative gene ontology terms significantly enriched for the h5UTRs according to criteria in Fig. 1C. A number of genes mapping to each term are also displayed (also see **Table S2**).

We next asked if h5UTRs are more likely to be discordant in their mRNA:protein expression levels, which would suggest post-transcriptional regulation. To do this, we used the GTex consortium transcriptomics and proteomics data which quantify mRNA and protein levels of each gene across 32 human tissues^28, 29^, to determine if genes with h5UTRs have a different distribution of per-gene cross-tissue correlations in mRNA versus protein levels compared to genes with similarly sized, non-conserved (defined as no overlap with LOD≥500 PhastCons elements) 5’UTRs. For 181 h5UTR genes, both RNA and protein expression was detectable in at least 10 tissues and the h5UTRs were annotated in both human and mouse RefSeq databases. Compared to all genes or to size-matched non-conserved controls, we observe significantly lower (Wilcoxon rank-sum test p=0.0013, p=0.0017 respectively) cross-tissue correlations for h5UTR genes (**Fig. 1B**). We also compared cross-tissue correlations of h5UTR genes with RNA variance-matched non-conserved controls to eliminate a model in which h5UTRs impact the correlations only through a different dynamic range of variation in RNA expression. The correlations were still lower for the h5UTR group (p=0.03) (**Fig. S1B**). In summary, protein levels of h5UTR genes, as a group, are more difficult to predict with RNA levels alone than those of non-conserved 5’UTR genes, suggesting that extreme sequence conservation in the 5’UTR may be due to tissue-specific post-transcriptional control.

To describe the potential biological functions of genes with h5UTRs, we surveyed gene ontology (GO) terms enriched in the h5UTR gene set. This survey identified 1037 significant GO terms at FDR≤0.05. To more stringently select the most informative set of GO terms amongst the 1037, we further used neighborhood-weighted scores that account for the overlaps among similar GO terms and also required a minimum enrichment ratio (**Fig. 1C**). This analysis resulted in a final list of 226 terms which we clustered into 22 groups by semantic similarity. To ensure the specificity of the enrichment, we also analyzed a length-matched set of non-conserved 5’UTR genes using the same procedure and threshold, which did not yield any enriched term (**Fig. S1C**). Thus, there is a strong and specific trend for h5UTRs to be found in genes driving distinct biological functions. Most strikingly, h5UTR GO terms highlighted genes critical for vertebrate embryonic developmental processes (**Fig. 1D, Table S2**). For example, h5UTR genes are annotated for morphogenesis of major tissues and organs including the circulatory, nervous, respiratory, reproductive, and skeletal systems. The nervous system is the largest among these, with a multitude of subcategories covering behavioral, cellular, and molecular processes, for example, learning, axon guidance, or synaptic transmissions. Genes that are part of signaling pathways involving the molecules Wnt, retinoic acid, GABA, Fgf, activin, BMP, Pdgf, Notch, Vegf, Hedgehog or Semaphorins are also abundantly present. We also note the genes involved in transcriptional programming such as chromatin remodeling and histone acetylation. Lastly, while UTRs are often neglected from mapping of genetic variants and their function, we nevertheless intersected known disease-associated variants with h5UTRs and found 5 potentially interesting associations (**Table S3**) ^30^. Overall, these annotation enrichments suggest that h5UTRs may play an important role in the post-transcriptional control of core embryonic developmental regulators.

### Hyperconserved 5’UTRs impact translation efficiency

We next sought to experimentally address whether the h5UTRs could impact the translational efficiency of mRNAs. In particular, we chose five candidates - Chrdl1, Gdf5, Dlx1, Sema3a, and Zfx - that function in contexts where spatiotemporal expression patterns are important for normal embryonic development. Chrdl1 is a member of the chordin family of BMP antagonists with numerous functional roles in cell differentiation and synapse plasticity, implicated in multiple neurological disorders^31–36^. Gdf5 is a TGF beta family protein with roles in skeletal and nervous system development^37–40^. Dlx1 is a homeobox transcription factor that has critical roles in craniofacial patterning, as well as in the differentiation and survival of neurons in the brain^41, 42^. Sema3a is a semaphorin family protein that is secreted and functions as a guidance cue for axons and vasculatures^43–47^. Zfx is a X-linked transcription factor protein that regulates self-renewal of embryonic and hematopoietic stem cells^48^.

To examine the contribution of h5UTRs, we introduced deletions into the 5’UTRs of Chrdl1, Gdf5, Dlx1, Sema3a, and Zfx using pairs of CRISPR/Cas9 sgRNAs targeting segments ranging between 50 to 200nt within the HCEs (**Fig. S2A-E**, **Table S4**). Deletions for four genes (Chrdl1, Dlx1, Sema3a, Zfx) were made in mESCs. Zfx transcript expression is detectable in mESCs natively, while for Chrdl1, Dlx1, and Sema3a, we can induce their transcript expression by treating mESCs with retinoic acid to promote differentiation (**Fig. S2F**). Because Gdf5 expression was not detectable in mESCs or RA-induced differentiated cells, the deletion for Gdf5 was made in 3T3 cells, where it was expressed. Next, we performed polysome profiling to compare the translation efficiencies (TE) of wild-type versus the ΔHCE mutants. Polysome profiling allows quantification of translation efficiencies independent from effects on transcript levels. We fractionated cycloheximide-treated cell lysates from wild-type and mutant cells on a sucrose gradient to separate mRNAs based on their ribosome occupancy **(Fig. 2A)**. Global translation levels displayed no difference between the wild-types and CRISPR/Cas9-mediated HCE knockout cells as observed by their total polysome traces **(Fig. S2G-K)**. We next quantified the distribution of individual candidate mRNA species across the fractions. The mRNAs that are more highly translated are expected to be present in heavier polysomes as they are bound by more ribosomes. Interestingly, we observe that for all five candidates tested, the deletion mutants exhibited a shift in the distribution of the targeted mRNA species from the heavier polysomes into the lighter polysomes, indicating a decrease in TE **(Fig. 2B-F)**. These findings suggest that h5UTRs may frequently harbor uncharacterized, additional cis-enhancers of translation initiation.

**Figure 2.**
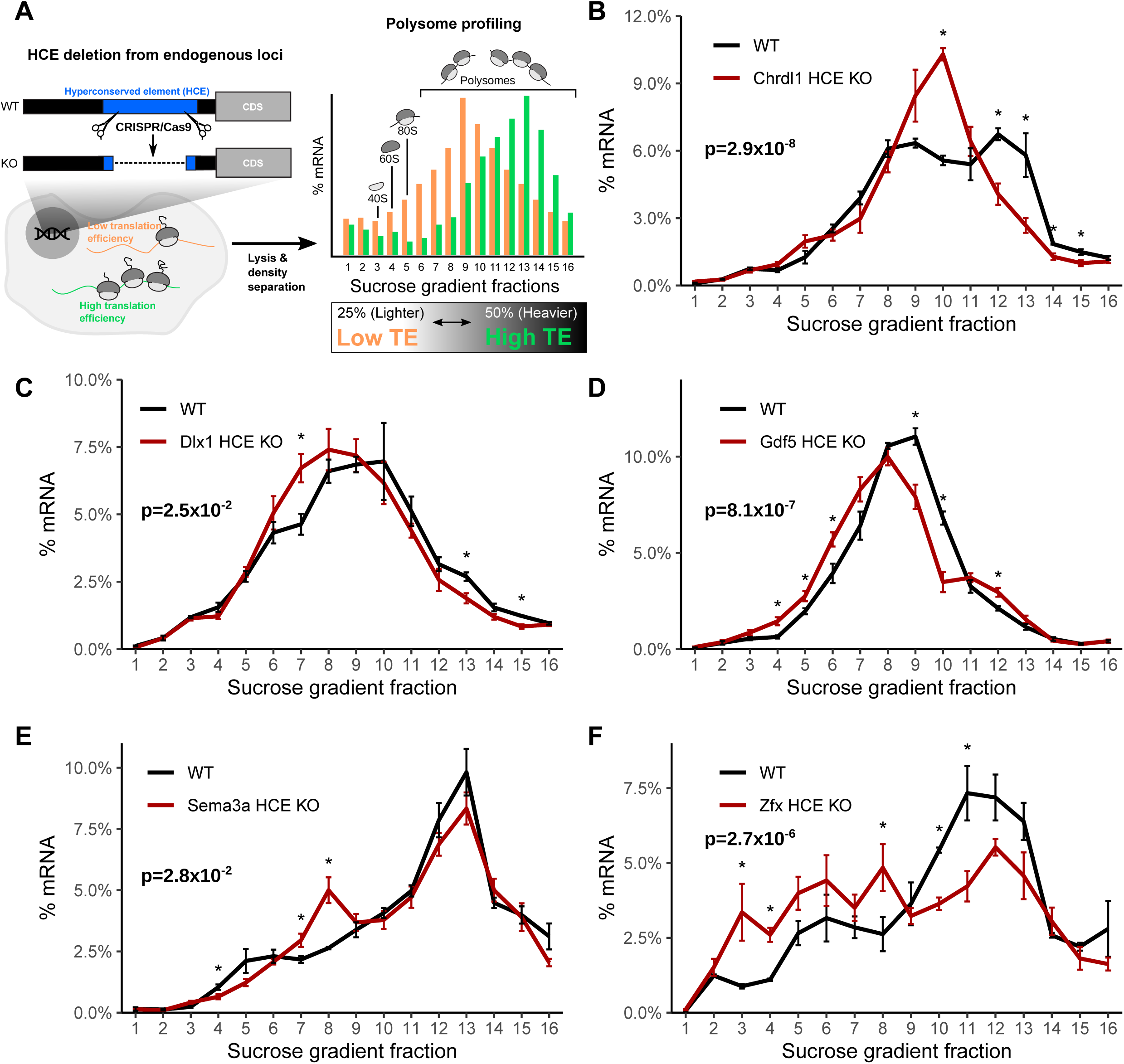
**Hyperconserved 5’UTRs impact translation efficiency** A) Schematic of experimental design for testing the impact of hyperconserved 5’UTRs on translation of coding genes. Shift in the distribution of the mRNAs across sucrose gradient fractions towards the right (heavier polysomes) indicates more average ribosome loading and higher translation efficiency (TE), while shift towards the left indicates lower translation efficiency. B) - F) Polysome profiles of wild-type versus hyperconserved element (HCE) knockout cells. Distribution of mRNAs across sucrose gradient fractions are plotted. Y-axis plots the mean percent mRNA. Error bars indicate standard error. Asterisk indicates t-test p≤0.05 for each fraction between the knockout and the wild-type. Indicated p-value (p_f_) is calculated by Fisher’s method across all fractions.

### Non-canonical translation initiation enhancer elements are a feature of hyperconserved 5’UTRs

A large fraction of the transcriptome is translated mainly via cap-dependent translation initiation mediated by the interaction of mRNA 5’ cap with eIF4E, the major cap binding protein. However, within specific genes, there has been growing evidence for the importance of less understood, alternative mechanisms of initiation independent of the cap-eIF4E interaction that have the potential for enhancing transcript-specific regulation of gene expression. For example, other eukaryotic translation initiation factors such as eIF3 or eIF4G components, or select ribosomal proteins of the ribosome, can interact with specific sequences in mRNAs to promote translation^26, 49, 50^. In addition, it has been estimated that 5-10% of cellular mRNAs may have the capacity to undergo cap-independent translation^51, 52^. However, it remains largely unknown what, if any, distinctive sequence or structural features predict and are predicted by such non-canonical translational enhancers. The HCE of Hoxa9 contains a functional RNA element previously shown to direct translation initiation in a cap-independent manner that is required for spatiotemporal control of Hox protein expression in the developing embryo^26^. Therefore, we asked whether other h5UTRs can similarly activate non-canonical translation initiation.

To test this hypothesis, we performed a large-scale reporter assay to measure the levels of non-canonical translation initiation from the h5UTRs. We synthesized and cloned a library of 253 full-length h5UTRs into a bicistronic reporter construct. In this system, two reporter genes, Renilla and Firefly luciferase, are transcribed through a SV40 promoter as one mRNA. The first cistron, Renilla luciferase, is positioned immediately downstream of the promoter and is translated by cap-dependent translation. The second cistron, Firefly luciferase, is positioned downstream of Renilla luciferase and can only be efficiently translated if the intercistronic inserted sequence enhances non-canonical translation initiation. The ratio of the two reporter luciferase activities provides a quantitative measurement of this activity. To perform the reporter assays, we initially selected the 10T1/2 cell line, a mesodermal cell line, as a pilot cell type and further expanded our analysis to include mESCs, neural stem cells (NSC), embryoid bodies, neurons, and primary cultures of limb bud mesenchyme to better represent a repertoire of lineages and differentiation trajectories in the developing embryo and capture instances of cell type-specific translation control.

We noticed two groups of reporter activities distributed in a bimodal distribution for each of the cell types (**Fig. S3A**). Within the higher group, we found the three positive controls we included in the reporter assays which all promote cap-independent translation - hepatitis C virus (HCV) internal ribosome entry site (IRES), encephalomyocarditis virus (EMCV) IRES, and the Hoxa9 h5UTR. The “empty” negative control reporter activity is found in the lower group near its median. Thus, the lower component appears to represent the background noise level present in our reporter assays. Using mixture modeling of the bimodal distribution, we estimated the false discovery rate for each tested h5UTR as the probability that the reporter activity of the tested h5’UTR could have come from the lower noise group. Using the maximum reporter activity across all six assayed cell types, we estimated that the proportion of the tested h5UTRs with non-canonical initiation activity is 33%. At 10% FDR, we are able to identify 90 h5UTRs with high non-canonical translation activity in at least one cell type (**Fig. 3A, Table S5)**. Of the 90 5’UTRs with non-canonical translation activity, two are previously known to the literature^53, 54^. The five genes (Chrdl1, Gdf5, Dlx1, Sema3a, Zfx) for which we demonstrated evidence of translational enhancers (Fig. 2B-F) fall in this class of 5’UTRs promoting non-canonical translation initiation. Strikingly, we observe that for 36 5’UTRs, their reporter activities are significantly variable across different primary cell types that we tested (**Fig. 3B**). To additionally examine whether cell-type specific non-canonical translation may be relevant in an endogenous context, we compared the polysome profiles of h5UTRs that show increased bicistronic reporter activity in NSCs relative to mESCs. Of the 9 h5UTRs analyzed, 5 (Gpbp1, Ppp2r5e, Pten, Trpm7, Senp3) showed shifts that indicated significant increase in translation in NSCs over mESCs, despite lower global translation in NSCs compared to mESCs (**Fig. S3B-L**). These results suggest that non-canonical translation initiation mediated by h5UTRs could be controlled in a highly regulatable fashion across different cell types to differentially control post-transcriptional gene expression.

**Figure 3.**
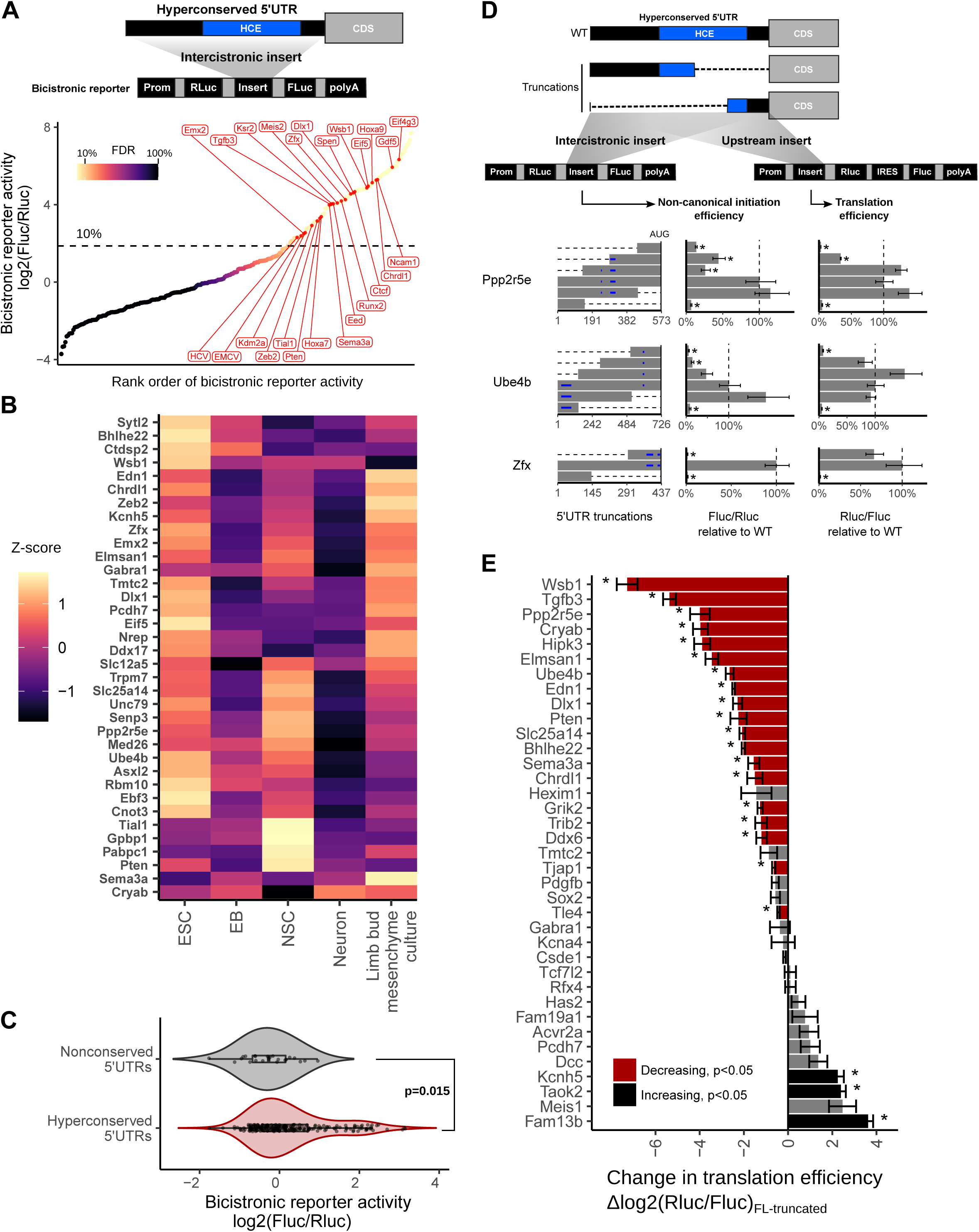
**Non-canonical translation initiation enhancer elements are a feature of hyperconserved 5’UTRs** A) Measurement of non-canonical translation initiation activity from 253 hyperconserved 5’UTRs by bicistronic reporter assay. Each dot is a 5’UTR, where x-axis is the maximum luciferase reporter ratio across six different cell types and y-axis is the rank of the reporter ratio from low to high. The skewing is reflective of the bimodal distribution of the activities (see also **Fig. S3A**), and color of the dot indicates estimated proportion of false positives based on mixture modeling of two Gaussian distributions. Dashed line indicates the reporter ratio above which 10% of the hits are expected to be false positives. Genes labeled in red: HCV and EMCV are positive control viral IRES; others are select h5UTRs with annotated biological functions in embryonic development. B) Heatmap of non-canonical translation initiation activity for 36 significantly varying h5UTRs across five indicated cell types (F-test, FDR≤0.05). The color shows row z-scaled reporter activities. The 5’UTRs are ordered by clustering similar reporter activity patterns across cell types. C) Violin plot of bicistronic reporter activities from hyperconserved and non-conserved 5’UTRs in 10T1/2 cells. p indicates Wilcoxon rank sum test p-value. D) The effect of various truncations of the h5UTRs on non-canonical initiation and total translation efficiency. Also see **Fig. S3M**. Left: positions of truncations. Dashed lines indicate truncations and bars indicate the remaining sequences. Blue horizontal lines within bars indicate uORFs; green and red lines within bars indicate in-frame and out-of-frame uAUGs, respectively. Middle: non-canonical initiation efficiency. Right: total translation efficiency. X-axis indicates the mean luciferase reporter ratios relative to the wild-type. Error bars indicate standard error. Dashed line marks the reporter ratio for the wild-type 5’UTR. N=4. Asterisk indicates t-test p≤0.05 for each truncation mutant versus the wild-type. E) Comparison of translational activities between the full-length h5UTR versus the only first 300nt of the h5UTR. 38 different pairs are tested. Error bars indicate standard error. Bars colored in red indicate significantly reduced translation in the shorter, truncated 300nt fragment; black indicates significant increase (t-test p≤0.05, marked by asterisk).

We also determined that the higher reporter ratios are not due to cryptic transcriptional and splicing effects, since the ratios of the mRNA levels of the two luciferase genes measured by qPCR in transfected cells are not skewed or correlated with ratios of the two luciferase reporter activities (**Fig. S3L**). In addition, we selected 23 non-conserved mouse 5’UTR sequences and tested their activities in 10T1/2 cells. Comparing these to the reporter activities of the h5UTRs in the same 10T1/2 cell type, the distribution of non-conserved 5’UTRs are unimodal near the lower noise component of the mixture observed for the conserved set (Wilcoxon rank sum test p=0.015, **Fig. 3C**). This result additionally supports our estimate of the proportion of the tested h5UTRs with non-canonical initiation activity. It further indicates that the extreme conservation in the 5’UTRs enriches for non-canonical translation initiation, suggesting their predictive value in identifying such elements genome-wide.

As all eukaryotic mRNAs are capped, the predominant mode of translation initiation is thought to be cap-dependent. It has been often argued that cap-independent translation typically makes only minimal contribution to overall translation efficiency, except under conditions of cellular stress, mitosis, apoptosis or other conditions during which canonical cap-dependent translation is globally reduced^55–57^. To understand the contribution of the non-canonical translational activation by the h5UTRs in the native context where cap-dependent translation is also active, we performed two different reporter assays with a series of truncated h5UTRs. The first reporter assay was the bicistronic reporter assay as described above. In the second reporter assay, the same truncation variants were positioned in the upstream position before the Renilla luciferase, just as it would be oriented in the endogenous capped monocistronic mRNA. This experiment measures total translational levels for truncated h5UTRs in the native context where cap-dependent translation is active. For 9 out of 11 h5UTR truncations in the two reporters transfected to 10T1/2 cells, we observed that at least one truncation significantly reduced non-canonical initiation as well as the total translational levels **(Fig. 3D, Fig. S3M**). The trend for truncations to frequently reduce total translation activity is notable, since cap-dependent translation initiation typically increases in efficiency when the 5’UTR is shortened. We asked if this is more generally true in a larger set of 38 h5UTRs, by comparing the total translation directed by the full-length versus only the first 300 nucleotides of the h5UTR without a large proportion of the HCE in each h5UTR. 20 out of 38 decreases significantly in the shorter truncated 5’UTR relative to the full-length, while only 5 significantly increases (**Fig. 3E**). The relatively low translation in truncation variants indicates that the removed sequences are typically acting to increase translation. In contrast, truncating long, non-conserved 5’UTRs do not show the same trend for decreased translation (**Fig. S3N, S3O**). Furthemore, there is no correlation between the change in the density of upstream AUGs and change in reporter activities, indicating that the decreased translation in truncated h5UTRs is unlikely to be due to an unexpected increase in the repressive contributions from upstream translation start sites (**Fig. S3P**). Taken together, non-canonical translation enhancer elements in h5UTRs widely impact total translation efficiency in physiological cellular conditions, suggesting that h5UTR genes may be translated via more specialized initiation mechanisms that utilize evolutionarily constrained, sequence-specific cis-regulatory features.

### Cellular remodeling of the RNA structural ensemble in hyperconserved 5’UTRs

Higher order structures are inherent features of RNA molecules that underpin their biochemical function not only in non-coding RNAs like tRNAs or rRNAs but also in mRNAs where they can encode regulatory function, for example, by mediating ligand or protein binding with specificity that cannot be achieved with primary sequence alone. One powerful computational approach for finding functional RNA structures in the genome has been the covariation model in which the evolutionary alignment of RNA sequences across different species is used to identify similar patterns of variation that support conservation of base pairs in a low free energy structure^58^. However, the majority of previous computational predictions of RNA structures in vertebrates occur in “moderately” conserved regions of the genome and miss the HCEs^59–62^. This is because covariation signal is in nucleotide substitutions, which requires conservation sufficient for alignment but also variation that would enable statistical power to measure covariation. Since extreme conservation limits the extent to which covariation signals can be informative, addressing this question currently requires additional experimental data.

One key conclusion from previous transcriptome-wide in vivo RNA chemical probing experiments is that mRNAs are globally “unfolded” inside eukaryotic cells^63–65^. Inactivation of ATP-dependent RNA helicase unwinding activity by depletion of ATP led to the cellular RNAs becoming more structured^63^. At the mRNA 5’UTR, cap-dependent translation is known to require DExD/H RNA helicases such as eIF4A and DHX29 for scanning of highly structured 5’UTRs^66^. In addition, for many structured, non-coding RNAs in ribonucleoprotein complexes such as ribosomes or spliceosomes, RNA helicases frequently act to remodel the alternative or intermediate structures towards their functional form^67^. We postulated that specific regions of mRNA that display localized sensitivity in their structures to active cellular remodeling by RNA helicases could potentially lead us to functionally relevant structures within h5UTRs.

While previous genome-wide in vivo chemical probing experiments provided key global insights, they were limited in their statistical resolution on many mRNA regions due to the lack of high coverage necessary to resolve localized accessibility pattern difference between different conditions^63–65^. To obtain a high coverage accessibility data for a large number of h5UTRs, we performed a highly multiplexed adaptation of dimethyl sulfate (DMS) mutational profiling^68–70^. In DMS mutational profiling, adduct-induced reverse transcriptase misincorporation of dNTPs into cDNAs are used as the readout of DMS modification rates. This allows amplicon sequencing of targeted RNA regions of interest. We performed DMS treatment of mESCs under the conditions of no treatment or depletion of ATP using 2’-deoxy-D-glucose (2DG) and NaN_3_ (**Fig. 4A**). We aimed to profile 100 h5UTRs natively expressed in mESCs by designing a total of 384 250nt amplicons tiling across them in a multiplex PCR. Of these, 161 amplicons across 69 h5UTRs passed final quality control filters. Accessibility patterns under ATP depletion and no treatment are globally highly similar to each other overall, relative to the accessibility profiles generated from purified total RNA extracts which were denatured and re-folded in vitro (**Fig. 4B**). We developed a statistical framework to analyze differential accessibility across our replicated data, which identified 140 11nt windows over 20 h5UTRs that were significantly different between ATP depletion and no treatment at a 5% false discovery rate (FDR) (**Fig. 4C, Table S6**). This represents a set of loci that are alternatively structured as a consequence of ATP depletion based on their accessibility patterns across each nucleotide. One known source of RNA structure remodeling in the cell is ribosome unwinding of mRNAs during translation, and thus the presence of upstream open reading frames (uORF) may lead to differential accessibilities upon ATP depletion^71^. We tested whether differential accessibility windows (FDR≤0.05) are overrepresented in upstream AUGs or potential uORFs but did not observe significant enrichment for either case, arguing against this possibility (**Fig. S4A, S4B**). Together, these results suggest the frequent presence of secondary structures under active energy-dependent cellular remodeling within h5UTRs.

**Figure 4.**
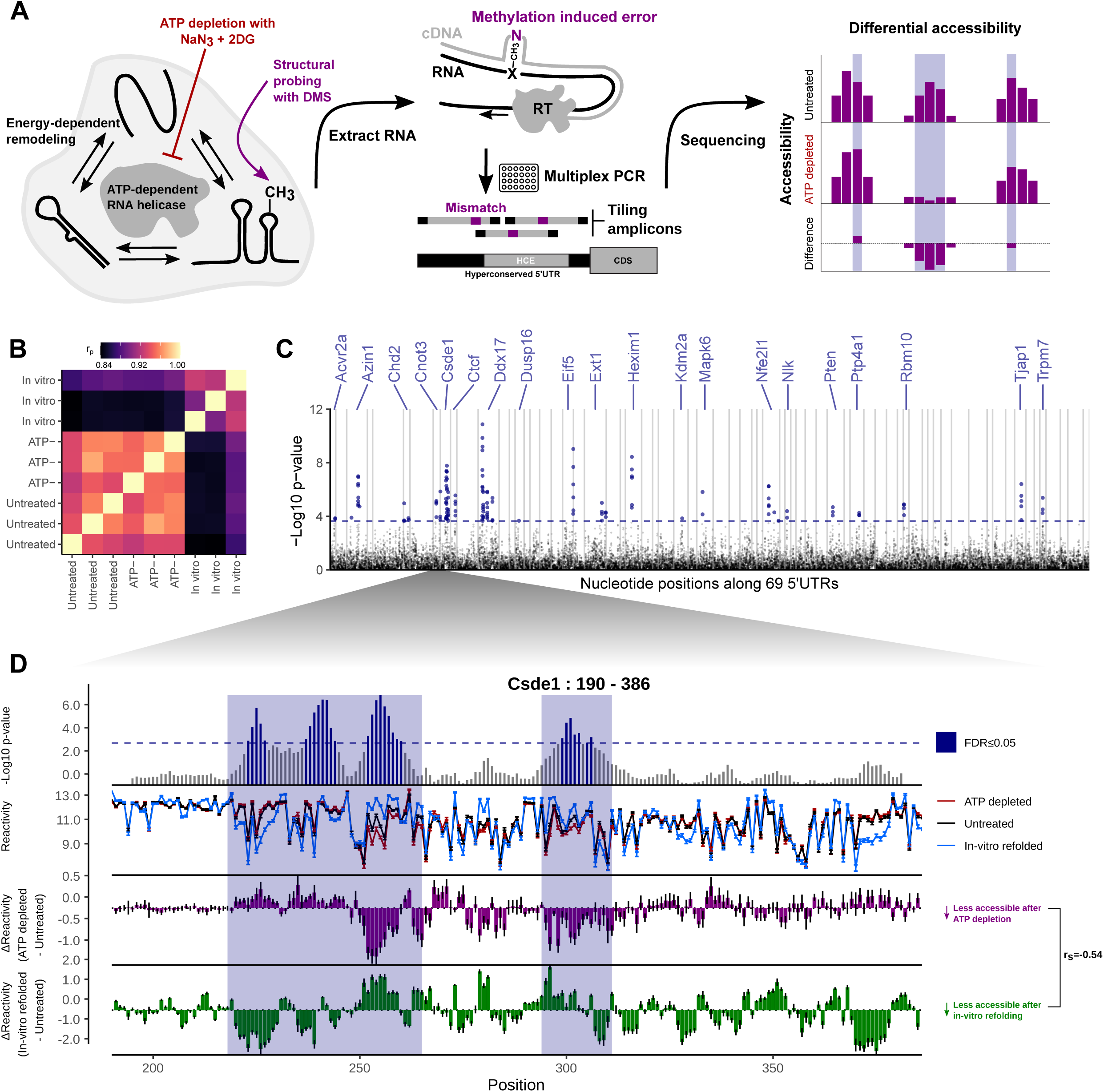
**Cellular remodeling of RNA structures in hyperconserved 5’UTRs** A) Schematic of identifying RNA structures under cellular remodeling in h5UTRs. Multiplexed, targeted DMS chemical probing of 69 h5UTRs inside cells from their endogenous mRNAs is performed following ATP depletion treatment to stop RNA helicase activity. B) Heatmap of correlation matrix across replicate samples for untreated, ATP depleted, and in vitro refolded samples (three each). The correlation values are calculated from a vector of normalized accessibility values for all nucleotides passing per-amplicon reproducibility cutoff. C) Manhattan plot of differential accessibility tests in 11nt overlapping windows across the 5’UTRs. Y-axis indicates -log10 KS-test p-value for each window along 69 5’UTRs in x-axis. Dashed line indicates the p-value cutoff at which permutation FDR is at 5%. D) Zoomed-in view of differential accessibilities along the Csde1 5’UTR in from positions 190 to 386. Top plot shows -log10 p-value for each window. Highlighted boxes mark significantly different windows, above the dashed line indicating 5% FDR. Middle plot shows differential accessibility on the y-axis, where greater than zero indicates increased accessibility upon ATP depletion and less than zero indicates decreased accessibility. Bottom plot shows differential accessibility for in cell versus in vitro refolded RNA.

To explore potential functional associations of the structural remodeling in h5UTRs, we intersected the list of 20 significant differential accessibility window-containing h5UTR genes with mammalian phenotype ontology. We found strong enrichment of terms that indicate essential early developmental gene function: 16 out of the 20 were annotated for either embryonic or neonatal lethality (**Table S7**). We also found known associations with human genetic diseases for 6 out of the 20 (**Table S8**). This suggests that the extremely conserved, structured RNA elements could be impacting post-transcriptional regulation of key developmental genes.

Among the most striking patterns of ATP-dependent differential accessibility observed is in the 5’UTR of Csde1 gene. The Csde1 gene, also known as upstream of N-ras (Unr), encodes a RNA binding protein (RBP) that contains five cold shock domains and regulates the translation and stability of its target mRNAs. Its molecular function in post-transcriptional regulation impacts a wide range of cellular processes including cell cycle, stem cell differentiation, apoptosis, and dosage compensation^72–78^. Furthermore, its expression levels are implicated in a variety of human diseases including Diamond-Blackfan anemia, autism spectrum disorders, and cancers^79–82^. We identified an approximate 150bp stretch from positions 215 to 365 encompassing a HCE that shows large scale accessibility changes upon ATP depletion with multiple statistically significant peaks (**Fig. 4D**). This suggests the possibility that active cellular remodeling of secondary structures in the Csde1 h5UTR may be functionally important for the precise regulation of its expression levels.

Notably, the accessibility changes observed in Csde1 h5UTR following ATP depletion are different from the changes observed between in cell and in vitro refolded RNA - there is even a slight negative correlation (r_s_=-0.54). Such discordance is also observed in a number of other h5UTRs with ATP-dependent differential accessibility (**Fig. S4B, Fig. S4C**). These patterns do not conform to the previously reported global trend in which there is a strong positive correlation between structures under ATP depletion and under in vitro conditions^63^. Thus, higher per-nucleotide signal-to-noise ratio data here indicate that for many h5UTRs, active remodeling by RNA helicases can be important for the formation of cellular structures distinct from those formed under in vitro conditions.

### In-cell mutate-and-map resolves functional alternative RNA structures encoded in the hyperconserved Csde1 5’UTR

As a paradigm to investigate cellular RNA structure and its remodeling in mRNA untranslated regions, we sought to further characterize the helicase-sensitive structures in Csde1 h5UTR. In particular, we took advantage of a modified mutate-and-map (M^2^) strategy^83^. In traditional M^2^, systematic mutagenesis of the RNA in question is coupled with chemical mapping, where separate accessibility profiles are generated for every mutated nucleotide. Effectively, this results in a two-dimensional matrix, allowing resolution of ambiguities typical in conventional chemical probing analysis where the one-dimensional data often cannot distinguish between conflicting structural models. Two different informative patterns are typically observed in M^2^ data. First are localized perturbations: here, the mutants do not disrupt the overall structure and increase the accessibility of only their interaction partners, thus revealing an approximate contact map **(Fig. 5A)**. Second are correlated global perturbations: the mutants produce large-scale accessibility changes across the RNA molecule in a correlated fashion, revealing the presence of multiple stable conformations that become more or less energetically favored as a result of the mutations (**Fig. 5B**).

**Figure 5.**
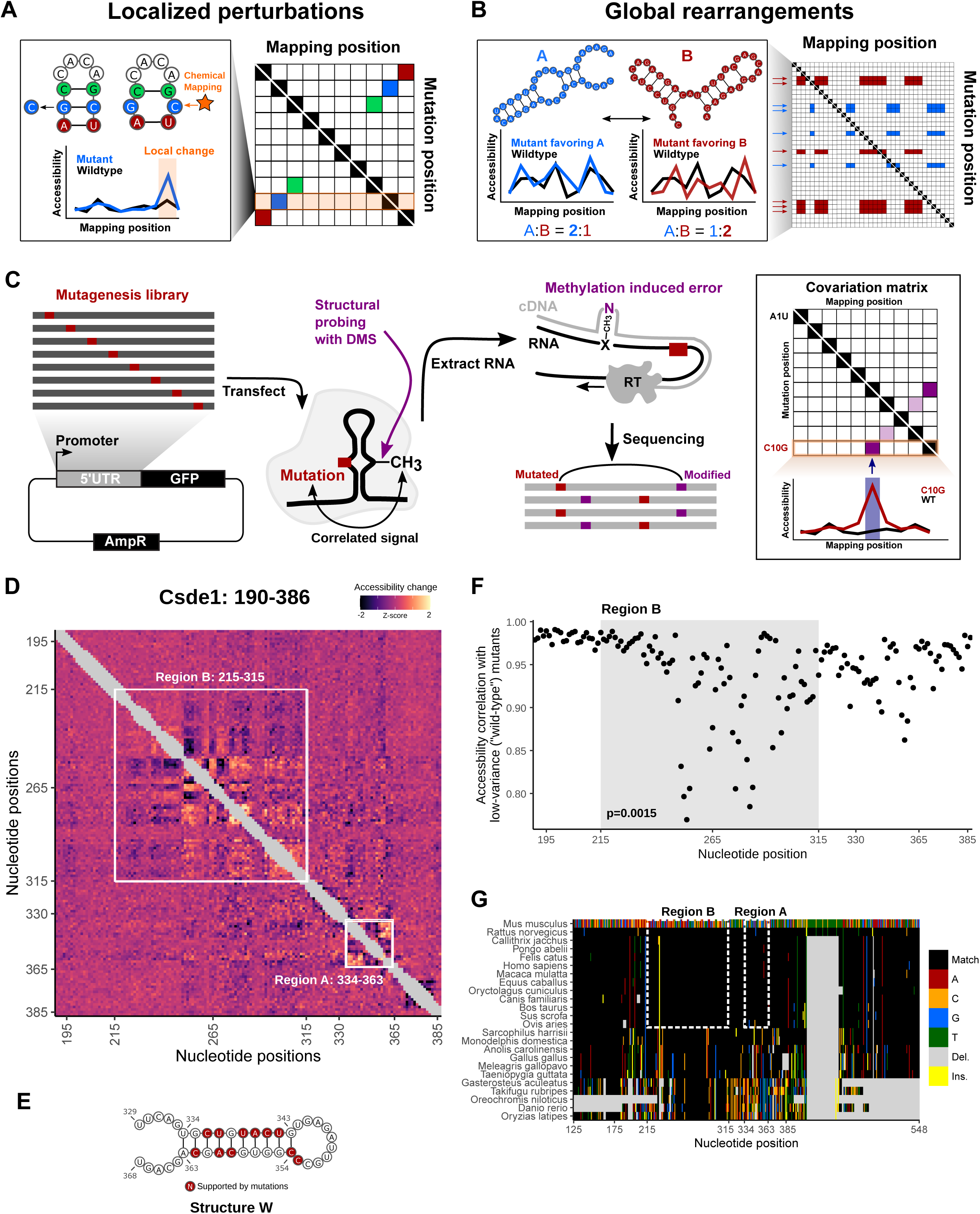
**icM^2^ reveals structured elements in the hyperconserved Csde1 5’UTR** A) Schematic of localized perturbation patterns that may be observed in M^2^ data. Here, the mutant does not disrupt the overall structure and “releases” its base pairing partner. This results in an increase of chemical accessibility signal at the interacting nucleotide. Systematic profiling of accessibilities by M^2^ results in an array of such mutant accessibility data into an approximate contact map. B) Schematic of global rearrangement patterns that may be observed in M^2^ data. Here, multiple conformations of the RNA molecule are present together in an ensemble at non-negligible relative proportions. Mutations can shift this balance, such that one structural state is favored over the other. In this case, M^2^ reveals large-scale accessibility perturbations across a longer stretch of the RNA molecule. Multiple mutations often impact the relative proportions in similar ways, which manifests as correlated arrays accessibility changes in M^2^ data matrix. C) Schematic of the icM^2^ method. Mutagenesis library of the target RNA of interest is first generated using error-prone PCR followed by cloning into an expression vector. The cells are transfected with the library and treated with DMS. Total RNAs are extracted. Read-through reverse transcription encodes DMS-modified nucleotides as mutations on the cDNA, which are read out by high-throughput sequencing. Correlated mutations in sequencing reads are then quantified and the resultant covariation matrix is analyzed for signature perturbation patterns. D) Heatmap of icM^2^ accessibility matrix for Csde1 5’UTR from position 190 to 386. For each row, the chemical mapping profile of a single-nucleotide variant of the RNA is plotted across the columns, where the colors indicate z-scaled accessibility change values from the wild-type RNA. 1D data from each mutant are vertically stacked to display a 2D matrix. White boxes mark the two regions (A: positions 334-363 and B: positions 215-315) that display strong perturbation signals that reveal their structures. E) A structure model (structure W) of region A. Bases colored in red indicate mutations with accessibility changes observed in icM^2^ data that are consistent with the model. F) Scatter plot showing correlations of per-nucleotide accessibilities between each mutant versus the “wild-type” (wild-type accessibilities are not directly measured, but mean accessibilities of 10 lowest variable mutants are used as a close approximation) on the y-axis and nucleotide positions along the x-axis. p indicates Wilcoxon rank sum test p-value for the difference in distributions of correlations between region B versus other nucleotides. G) Multiple species alignment for Csde1 5’UTR from position 125 to 548. For each row, the sequence alignment of a species is plotted across the columns, where the colors indicate match/substitution/insertion/deletion at each nucleotide. The alignment positions are relative to the mouse sequence. The top row is the mouse alignment, colored separately from other rows as a reference to indicate the identity of the bases in each position in the multiple species alignment.

A major limitation of traditional M^2^ is that its application has only been possible for RNAs under in vitro conditions. The influence of cellular constituents on RNA structure is critical and usually difficult to mimic by reconstitution: the microenvironments found in the cell differ in amounts of metal ions, metabolites, crowding agents, macromolecular interactors such as RBPs or helicase-mediated unwinding activities^84, 85^. Thus, we developed in-cell mutate-and-map (icM^2^), a powerful new methodology that enables the application of M^2^ strategy in the native cellular context (**Fig. 5C**). In icM^2^, the target sequence of interest is first mutagenized using error-prone PCR, cloned as a pool into an expression plasmid and transfected into cells. Following the treatment of transfected cells with DMS, total RNAs are extracted and subjected to read-through reverse transcription, where modified nucleotides are misincorporated as mutations on the cDNA that are amplified and sequenced. Correlated mutations in sequencing reads are then quantified, and the resultant covariation matrix is analyzed for signature perturbation patterns. icM^2^ is particularly suited for analysis of h5UTRs, as it directly addresses what RNA structural changes occur if each of the extremely conserved nucleotides are mutated during evolution.

We applied icM^2^ in three windows tiling across Csde1 5’UTR in mESCs. Strikingly, we observed strong perturbation signals in the ∼215-365 positions along the 5’UTR, where we had originally observed large differential accessibilities in response to elimination of RNA helicase unwinding activities (**Fig. 5D**). The visualization of the icM^2^ accessibility matrix immediately highlighted two subregions. The first region is around positions 334-363 (region A), where short range localized perturbations indicated the presence of a small stem loop motif. Here, the data corresponded well to the expected accessibility changes for the lowest free energy structure (structure W) predicted for the region (**Fig. 5E**). The second region is around positions 215-315 (region B), where correlated global perturbations across a long stretch of about 100 nucleotides indicated the presence of multiple conformations. Remarkably, these correlated global perturbations occur for almost every mutation across the 100nt stretch, revealing the strong sensitivity of the ensemble state to the precise sequence identity of each base. This is highlighted by plotting the correlation of per-nucleotide accessibilities between each mutant versus the “wild-type” (not directly measured but instead approximated from 10 lowest variable mutants), where an extended region of low correlations are observed (**Fig. 5F**, Wilcoxon rank-sum test p=0.0015 for mutants in region B versus other mutants). These results indicate that there are at least two alternative structures present in region B, whose relative proportions inside of the cell are affected by a mutation in almost any of the extremely conserved nucleotides. In addition, we observe the strongest conservation signal of the Csde1 5’UTR in region B (**Fig. 5G**). Upon closer examination of the alignments, we find that amongst placental mammals, in particular, there is a near-perfect sequence identity. The only variable nucleotide found in nature across the 100nt stretch occurs at position 229 (in mouse sequence coordinates), where there is a rodent-specific 2-bp deletion of AA and a marmoset-specific A to C substitution. These results suggest the presence of structured elements in the hyperconserved Csde1 5’UTR and a structural explanation for why such extreme conservation levels may be required at the level of RNA. Examining the conservation levels and ATP-dependent accessibility profiles across all other h5UTRs we DMS probed reveals that the average per-nucleotide conservation levels in significantly differential accessibility regions (FDR≤0.05) display exceedingly high conservation levels compared to the rest of the RNA (**Fig. S5A, S5B**). Thus, encoding of actively remodeled cellular RNA structures may be a broadly occurring phenomenon associated with the extreme conservation levels in h5UTRs.

We next asked what candidate structures might comprise and explain the observed alternative states of the ensemble in region B. Clustering the icM^2^ accessibility matrix revealed two groups of mutations with similar patterns of accessibility changes (**Fig. 6A**). This likely represents two separate structural states that dominate the ensemble as a result of the mutations, relative to the wild-type state balanced between the two. We used the average accessibility change profiles for the two clusters as two separate constraints for RNA folding. Constraining by cluster 1 average accessibility profile revealed a well defined conformation (structure X) disrupted by cluster 1 mutants in the helices and stabilized by cluster 2 mutants in the loops (**Fig. 6B**). Constraining by cluster 2 profile resulted in a higher entropy fold, which was nevertheless readily visualizable by two representative medoid conformations (structures Y and Z, **Fig. 6C, D**). To estimate the relative mixing ratios of these structures, we chose to apply RNA ensemble extraction from footprinting insights technique (also known as REEFFIT) which is an expectation-maximization algorithm that estimates an optimal linear combination of a priori list of structures to explain the observed M^2^ data^86^. For the wild-type sequence, REEFFIT yielded proportions of 67±9%:10±4%:23±9% for the representative structures X, Y, and Z, respectively (**Fig. 6B, C, D**). It also predicted how these proportions are expected to change across the individual mutants, adding quantitative estimates to our initially qualitative observations of alternative structural states. For example, cluster 1 mutants disrupt structure X to favor Y and Z, changing the relative proportions of X:Y:Z to 30%:13%:57% on average, while cluster 2 mutants act in the opposite direction, shifting it to 91%:6%:3% (**Fig. 6E**). We further discovered that the accessibility change profiles of cluster 1 mutants are closely correlated (r_s_=0.83) with our previous measurements of differential accessibilities produced under the condition of cellular ATP depletion. Conversely, the same ATP-depletion differential accessibilities exhibit negative correlation (r_s_=-0.8) with the accessibility change profiles of cluster 2 mutants (**Fig. 6F**). This observation suggests that elimination of RNA helicase unwinding activity, like cluster 1 mutants, decreases the proportion of structure X in the cell and does so to increase the fraction of the alternative structures Y and Z.

**Figure 6.**
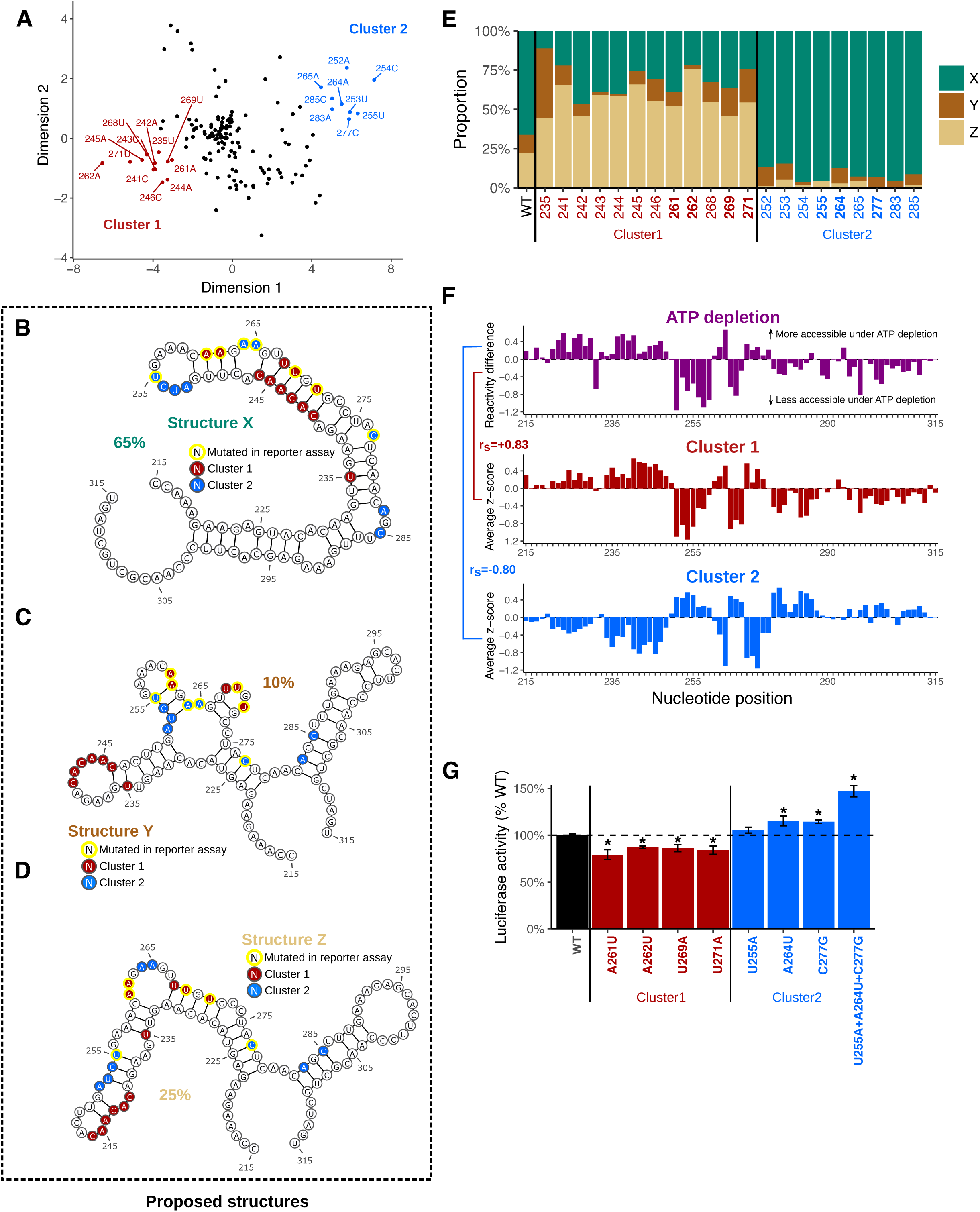
**Csde1 5’UTR tunes translation efficiency by encoding multiple alternative structures that are actively maintained by RNA helicases** A) Scatter plot showing dimensionality reduction of icM^2^ data matrix (positions 190-386) by classical multidimensional scaling (K=2). Dots indicate each single nucleotide variant of the RNA, where the colors/annotations mark the mutants grouped into two clusters, determined heuristically by visual inspection of the positions on the plot. B) - D) Structure models (X, Y, Z) for the alternative conformations in region B. Bases colored in red or blue indicate mutations with similar patterns of accessibility changes grouped into two clusters as shown in A). Yellow outline indicates mutants that are also tested for function in luciferase reporter assay (shown in Fig. 6G). Percentages indicate relative proportions of the three structures estimated by REEFFIT. E) REEFFIT estimates of the mixing proportions (relative to the maximal amount of change that can be observed by the single mutations) for structures X,Y, Z upon introduction of single nucleotide mutations. Y-axis indicates the stacked bars indicating proportions, along the variants from the two clusters in X-axis. F) Comparison of average accessibility changes for each of the two clusters with accessibility changes observed upon ATP depletion at region B. Top plot shows the reactivity differences upon ATP depletion, where greater than zero indicates increased accessibility upon ATP depletion and vice versa. Middle and bottom shows the cluster average accessibility change z-scores. Spearman correlation between cluster 1 and ATP depletion is 0.83 and -0.8 between cluster 2 and ATP depletion. G) The effect of shifting the relative balance of the alternative conformations at region B on translation. Y-axis shows changes in luciferase reporter activities compared to the wild-type sequence when the single variants affecting mixing proportions (shown in Fig. 6E) are introduced into the full-length Csde1 5’UTR upstream of the luciferase reporter. The bars and labels along x-axis are colored according to whether they are wild-type, cluster 1 mutants, or cluster 2 mutants. Dashed line indicates the wild-type luciferase reporter level. N=3. Asterisk indicates a significant difference between each mutant and wild-type t-test p≤0.05; U255A mutant is p=0.1).

Notably, structure X has multiple long stems (positions 232-282) - i.e. the helicase activity promotes a low free energy structure (−15.1kcal/mol for X versus -10.5kcal/mol for Y and -14.40kcal/mol for Z for the entire region; furthermore, ΔG of the motif in structure X at positions 232-282 is -9.30kcal/mol, while its counterparts for structures Y and Z at positions 223-279 are -7.70kcal/mol and -3.60kcal/mol, respectively). This potentially argues that under in-cell conditions, the other structures may be more locally stable, acting much like kinetic traps as often described for some non-coding RNAs where helicases act as chaperones^87^. It is formally possible for other direct contacts on the exact methylation sites of the nucleotides, such as a direct RBP interaction on the base-pairing face, to produce localized “footprints” on the accessibility profiles; however this would not drastically impact our model. Taken together, we propose three candidate conformations to account for our icM^2^ signal observed in region B of Csde1 5’UTR and hypothesize that the cell is actively spending energy to maintain the precise relative balance of these conformations in the cellular structural ensemble.

In our initial one-dimensional DMS profiling analysis of h5UTRs in mESCs, we had observed that the accessibility profiles of RNAs refolded in vitro were discordant from those of RNAs in cells (**Fig. 4D, S5A, S5B**). To further expand on these differences and actually compare in-cell vs in-vitro RNA structures, we also performed in vitro M^2^ on Csde1 h5UTR. We observed a strikingly different accessibility matrix which immediately hinted at well defined folds with longer range stems (**Fig. S6A**). In fact, our in vitro data clearly supported all helices seen in the minimum free energy structure that was blindly predicted from only the sequence (**Fig. S6B**). These results provide strong evidence that the energy landscape of RNA structures inside cells can be drastically different and highlight the importance of resolving flexible conformations that can occur uniquely under cellular conditions.

Lastly, we asked whether such a shift in the balance of structural conformations has a functional consequence on the translation of the downstream gene. To this end, we performed luciferase reporter assays with mutant Csde1 5’UTRs carrying a number of substitutions from each of the two clusters that are predicted to change the relative proportions. We selected four different nucleotide positions from cluster 1 and three from cluster 2, hypothesizing similar patterns of expression level changes may be observed among each cluster. We transfected capped and polyadenylated reporter mRNAs into mESCs, the same cell type in which we performed our structural analyses. We observed that all cluster 1 mutants decreased Firefly luciferase activities by 15-20% compared to the wild-type 5’UTR (**Fig. 6G)**. In contrast, cluster 2 mutants increased the reporter activities by 5-15%. When the three individual single mutations from cluster 2 are combined, the effect size is increased to about 50%. The dynamic range of final protein levels can thus be tuned according to the relative proportions of the multiple conformations along the RNA structural landscape of Csde1 5’UTR. Together, these results suggest that the exact proportions and properties of the RNA structural ensemble is a critical functional requirement under negative selection in hyperconserved vertebrate 5’UTRs.

## Discussion

Extreme sequence conservation, over stretches on the order of hundreds to thousands of nucleotides, has long been observed in non-coding regions of vertebrate genomes, yet our current functional knowledge of these elements falls short in explaining why and how such conservation levels exist. While genome-wide methods like ribosome profiling are allowing us to appreciate the widespread nature of transcript-specific translational regulation, a systematic understanding of cis-regulatory elements that explain the variation in translation efficiencies is still missing^88^. Here, we uncover a functional role for hyperconserved 5’UTRs in regulation of translation and report their unexpected enrichment in non-canonical initiation sites. Our results indicate that the evolution of cis-regulatory elements governing translation, particularly within those transcripts critical to the emergence of the most essential developmental features in vertebrate species, frequently circumvents canonical translation initiation while accommodating more specialized mechanisms. We speculate that there may potentially be many different types of unknown non-canonical mechanisms that are adopted by these 5’UTRs and that further investigations may identify new classes of RNA elements that are also utilized by other mRNAs in the genome. We suspect that a thorough analysis of mRNAs with h5UTRs may reveal a system of translational enhancers or repressors enabling an additional layer of spatio-temporal regulation of protein expression in development, akin to how large scale in vivo efforts on intergenic extreme conservation that have revealed a network of developmental transcriptional enhancers of mRNA expression. In addition, recent evidence indicates widespread alternative transcription start sites that lead to variable translation efficiency in mammalian cells^89, 90^. Notably, alternative 5’UTRs are also more frequently annotated for genes with h5UTRs than without (**Fig. S7**). It will thus also be interesting to explore whether and how expression patterns of h5UTRs may vary across tissues that may result in differential translatability of these mRNAs.

A crucial component of decoding cis-regulatory features at the level of RNAs is the determination of their higher order structures beyond the primary sequence. We found that cells precisely tune protein expression levels by remodeling the structural ensemble in the hyperconserved Csde1 5’UTR to maintain the relative proportions of multiple functional conformations. While icM^2^ revealed a highly dense array of mutations that disrupt such actively enforced balance of dynamic structures in the Csde1 5’UTR across a ∼100nt long stretch, the same mutations are negatively selected against in nature across vertebrate species, to the extent where it is almost completely invariant among mammalian genomes. This suggests that selective pressures for translational regulation can lead to extreme sequence constraints when an ensemble of multiple functional conformations must be encoded over a single stable structure to ensure a dynamic range of translational outputs. The observation that regions of h5UTRs under helicase-dependent structural remodeling in general display the highest conservation levels further suggests that a similar phenomenon could extend more broadly to other h5UTRs and may at least in part explain extreme conservation in 5’UTRs at the level of RNA. A number of disease-associated human single nucleotide polymorphisms that alter RNA structure, known as riboSNitches, have been suggested to influence dynamics of predicted RNA ensembles^91–94^. It will be thus interesting to see whether the intersection of human genetic variation across hyperconserved 5’UTRs and our icM^2^ methodology can identify novel riboSNitches with disease consequences.

Flexible structural states can potentially endow multiple functional states in regulatory elements that respond to environmental or cellular cues such as varying metabolite concentrations or RBP levels. Most current genome-wide efforts to identify functional structures, whether by covariation, thermodynamics, or experimental accessibility data, have focused on single stable conformations. Our results underscore the necessity of the ensemble perspective of RNA structure in understanding the cellular activities of regulatory RNAs and the potential utility of extreme conservation in detecting such dynamically structured elements in the untranslated regions of mRNAs. We envision that hyperconserved 5’UTRs will serve to catalyze the discovery of functional RNA structures in vertebrate genomes and to ultimately advance our broader understanding of post-transcriptional gene regulation in development, disease, and evolution.

## Author contributions

M.B., G.W.B, and E.S.C. conceived the project. M.B. supervised the project. L.J. and H.T. provided the GTEx data and critical feedback on its analysis. R.D. provided critical feedback on the development and analysis of icM2. E.S.C. carried out the large-scale reporter screens. G.W.B. performed all other experiments and data analysis. G.W.B and M.B. wrote the manuscript in consultation with all authors.

## Acknowledgements

We would like to thank the members of the Barna lab for constructive criticism of the manuscript. This work was supported by New York Stem Cell Foundation grant NYSCF-R-I36 (M.B.), NIH grant 1R01HD086634 (M.B.), Alfred P. Sloan Research Fellowship (M.B.), Pew Scholars Award (M.B.), Mallinckrodt Foundation Award (M.B.), Benchmark Stanford Graduate Fellowship (G.W.B.), Walter and Idun Berry Foundation (E.S.C.). M.B. is a New York Stem Cell Robertson Investigator.

## Figure legends

**Figure S1.**
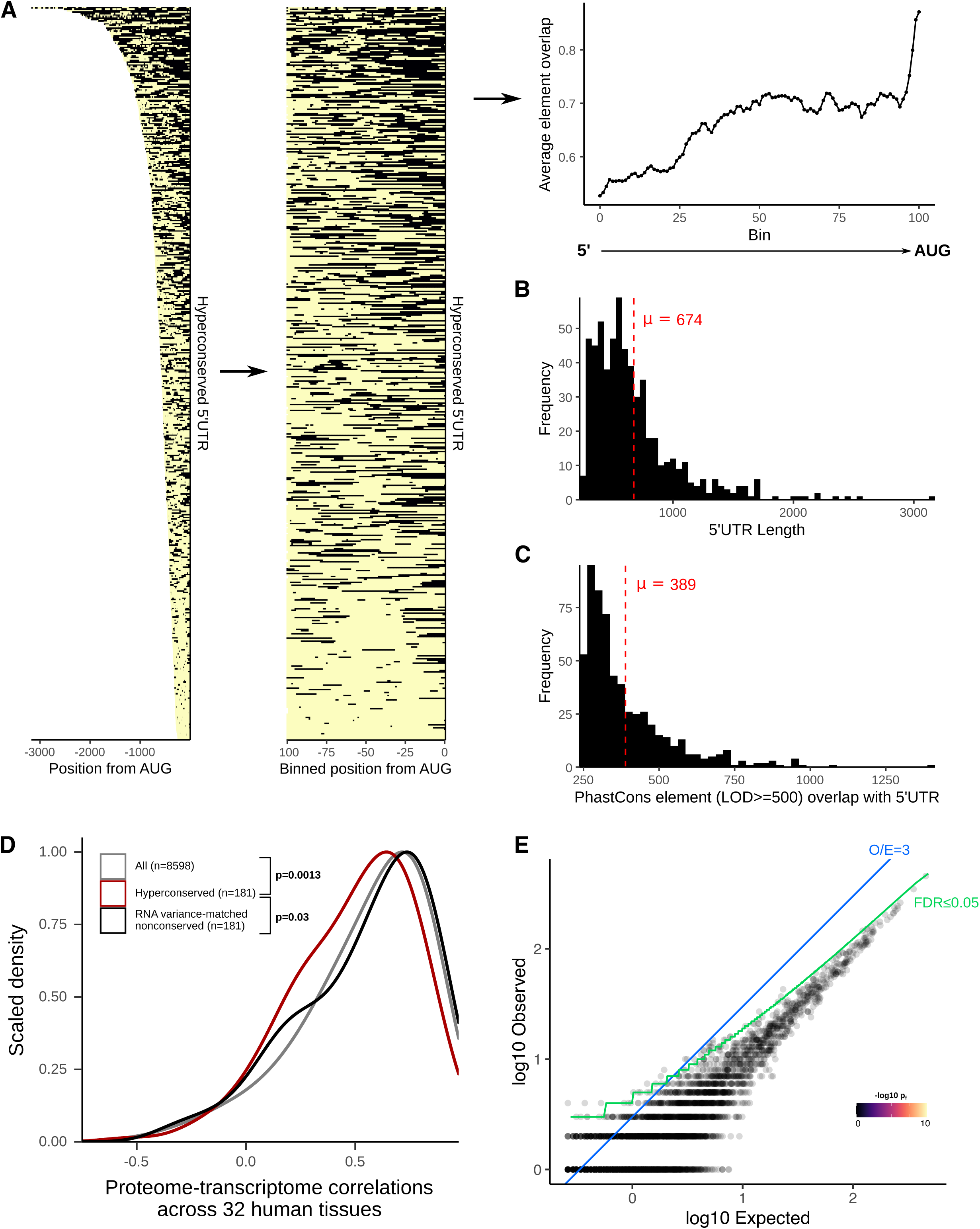
**Defining hyperconserved 5’UTRs in vertebrate genomes** A) Left: heatmap of the positions of LOD≥500 PhastCons elements in each h5UTR. Middle: heatmap of the relative positions (calculated in 100 bins across the h5UTRs) of the elements. Right: plot of average element overlap across the 100 bins to illustrate the positional preference. B) Histogram of the length of h5UTRs. Average length is 674nt. C) Histogram of the number of nucleotides overlap between LOD≥500 PhastCons elements and h5UTRs. Average overlap is 389nt. D) Distributions of cross-tissue transcriptome-proteome correlations for all genes, genes with h5UTRs, or genes with variance-matched non-conserved 5’UTRs. Indicated p-values are from Wilcoxon rank sum tests for cross-tissue correlation values between h5UTR genes and all genes or between h5UTR genes and variance-matched non-conserved controls. E) Scatter plot illustrating the lack of significant term enrichments for a size-matched set of non-conserved 5’ UTRs. X-axis and y-axis plots expected and the observed number of genes for each term. Blue dashed line indicates the minimum observed/expected ratio cutoff of 3. Green line indicates expected and observed counts where Fisher’s test p-value (p_f_) is estimated to have FDR=0.05. Neighbor-weighted test p-value (p_fw_) ≤0.05 is further used as an additional cutoff. The final set of enriched terms passing filter is colored by p_f_ and sized by p_fw_.

**Figure S2.**
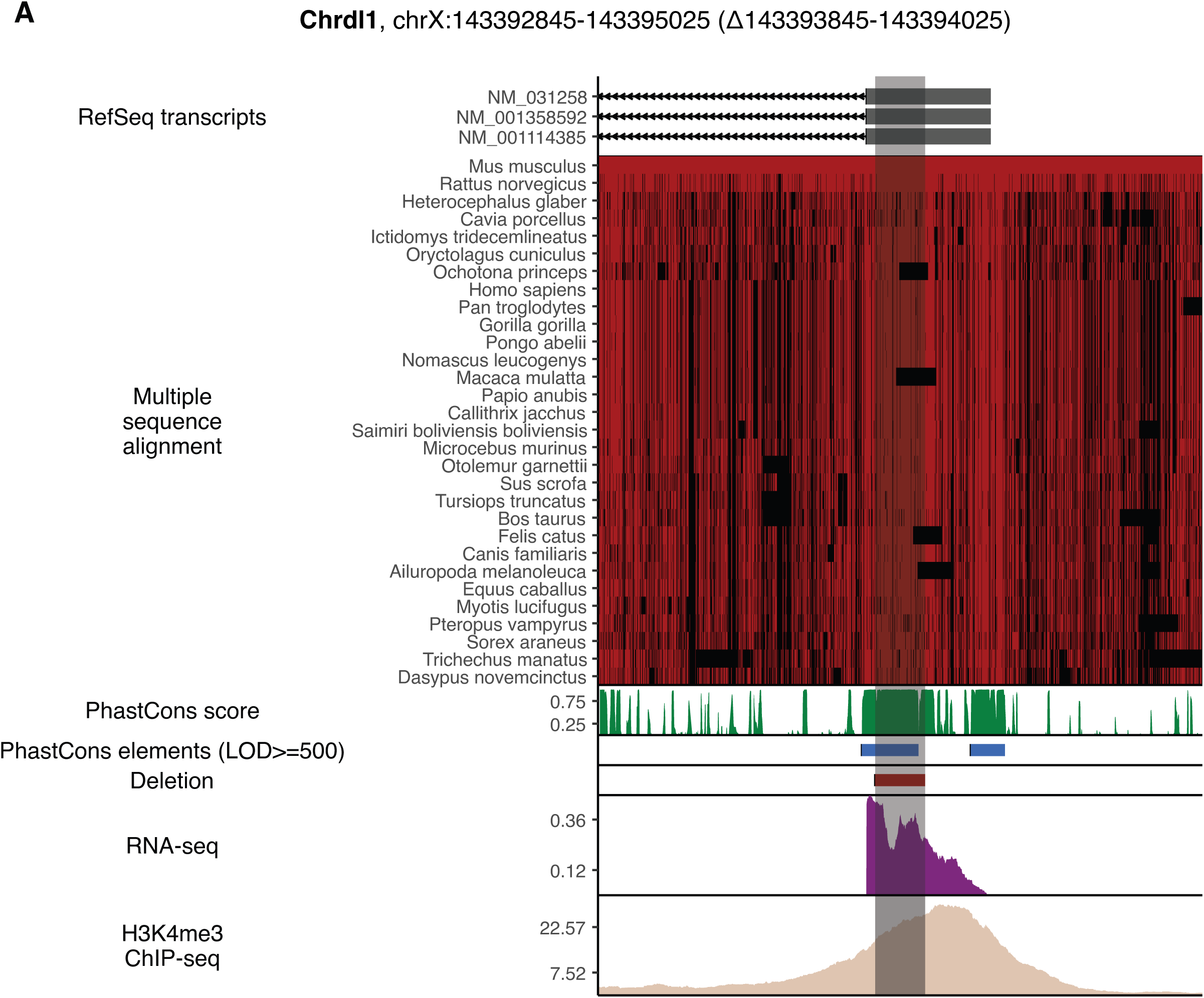
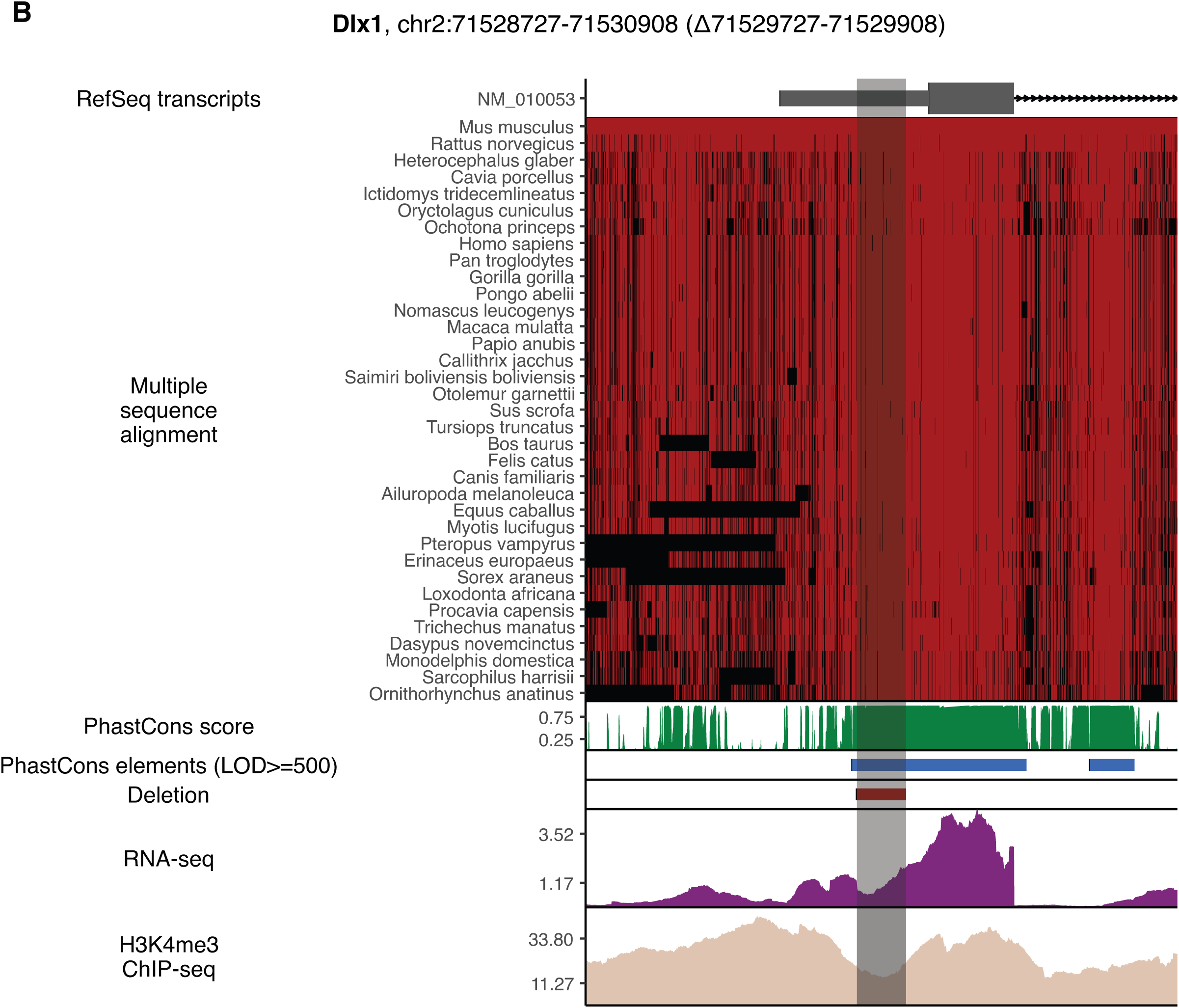
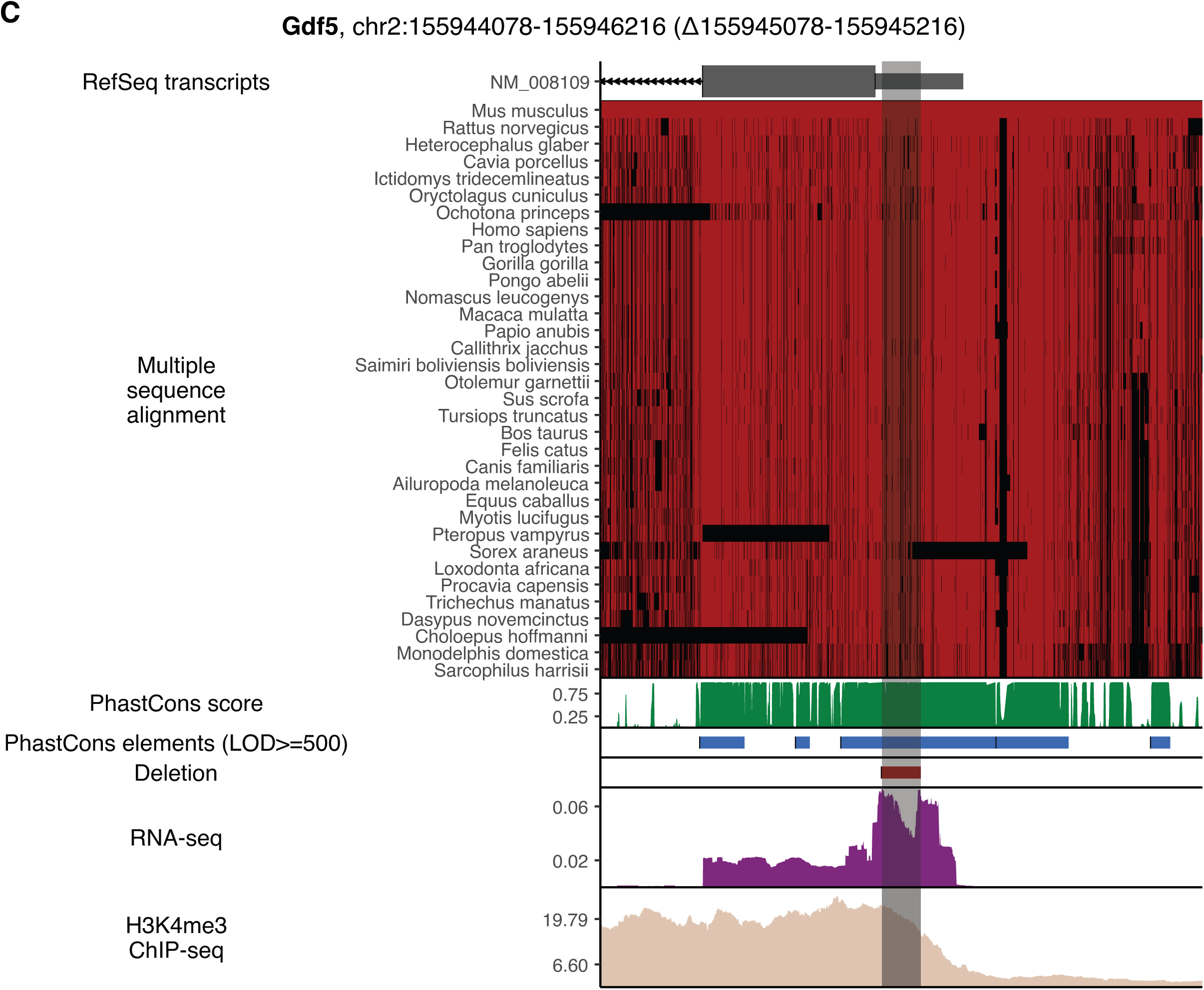
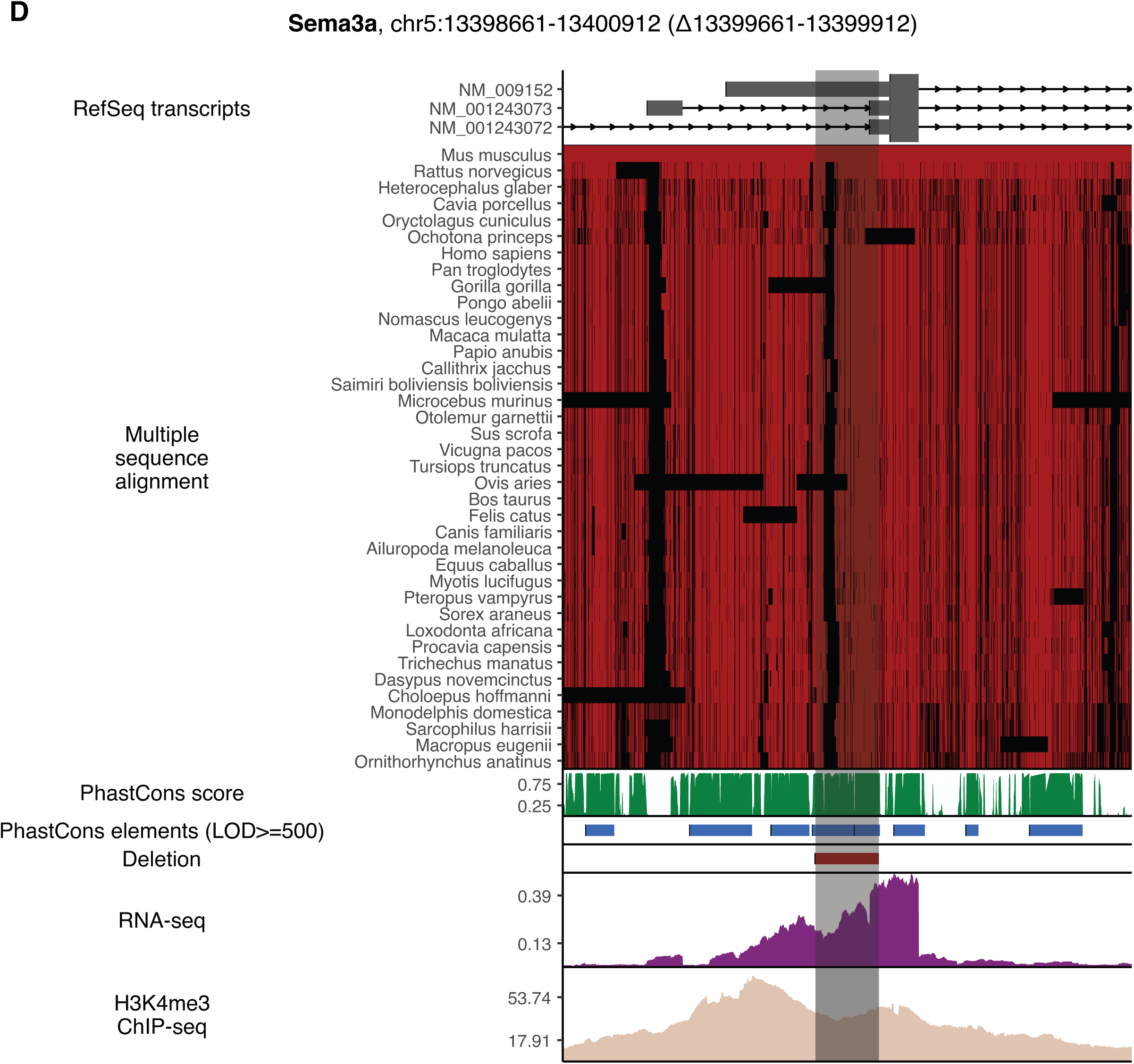
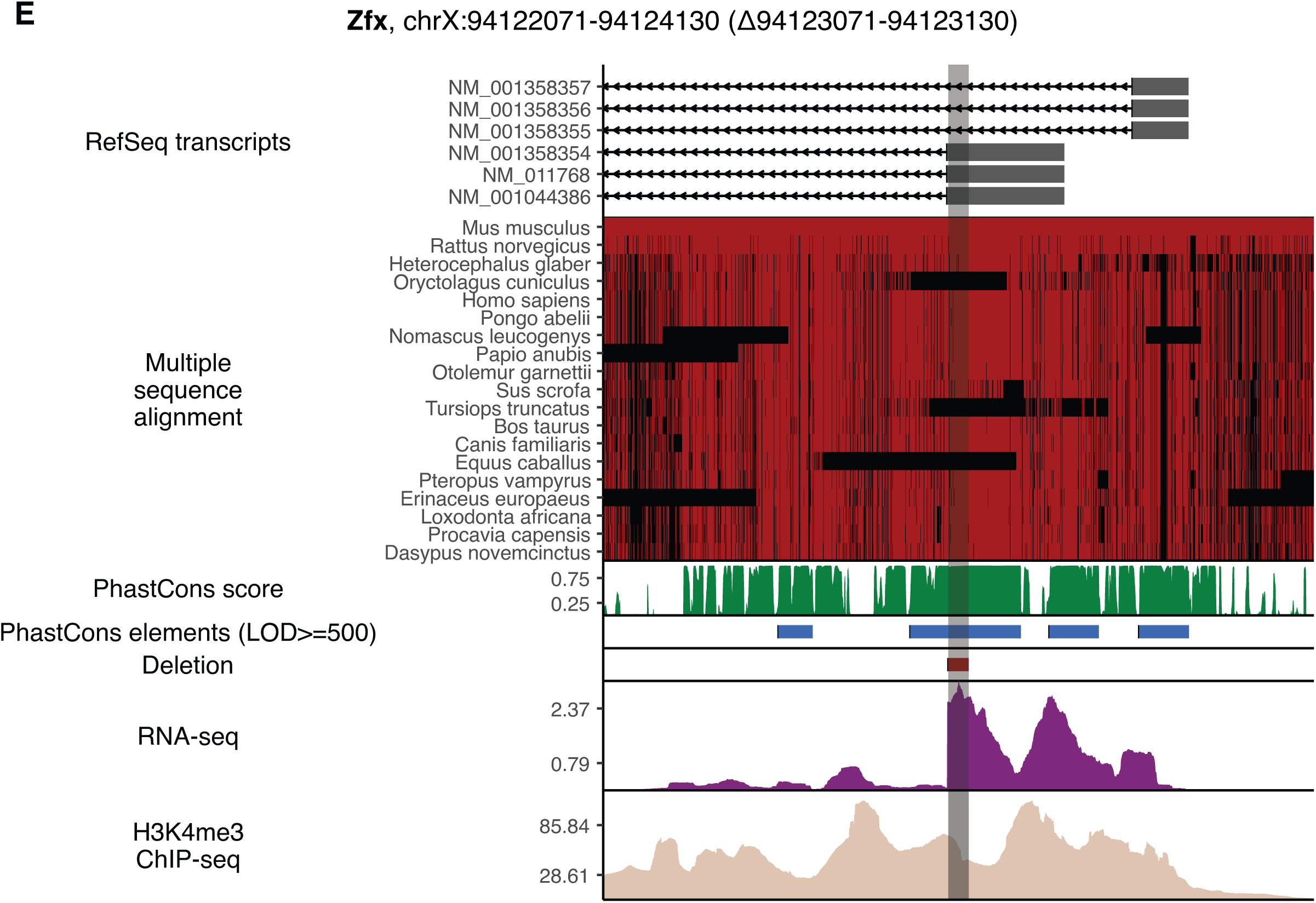
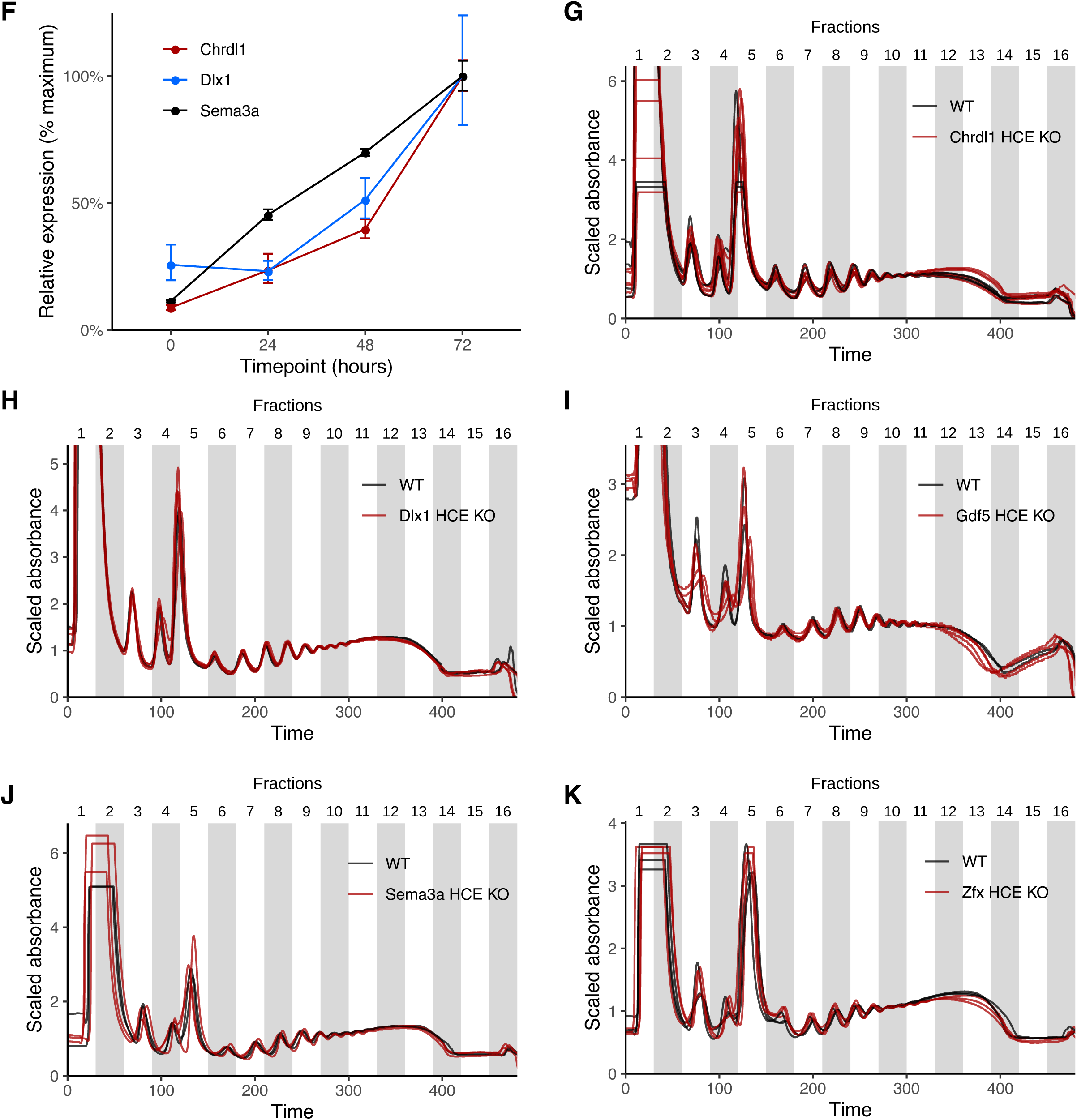
**Hyperconserved 5’UTRs impact translation efficiency** A) - E) Genome browser tracks illustrating the position, multiple sequence alignment, PhastCons scores, LOD≥500 PhastCons elements, and the location of the deletion. Aggregate (maximum signal across all datasets) RNA-seq and H3K4me3 ChIP-seq tracks are also shown. F) Induction of expression of Chrdl1, Dlx1, and Sema3a by retinoic acid treatment of mESCs. On the x-axis is the time after the addition of retinoic acid. Y-axis plots the mRNA expression level normalized to the maximum value for each gene. G) - K) Polysome traces of wild-type versus HCE knockout cells. Y-axis plots the relative A260 units and X-axis is the time along the fractionation.

**Figure S3.**
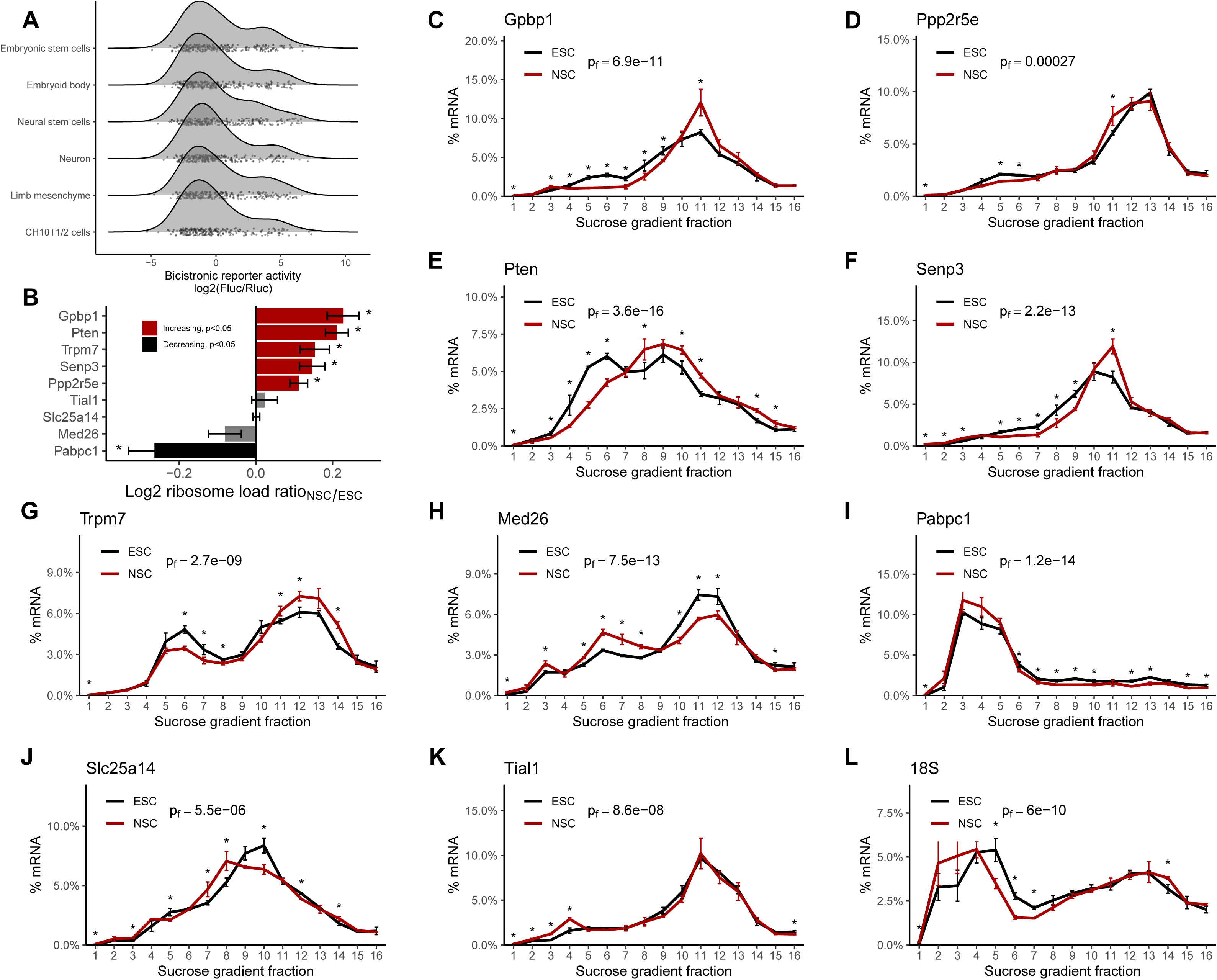
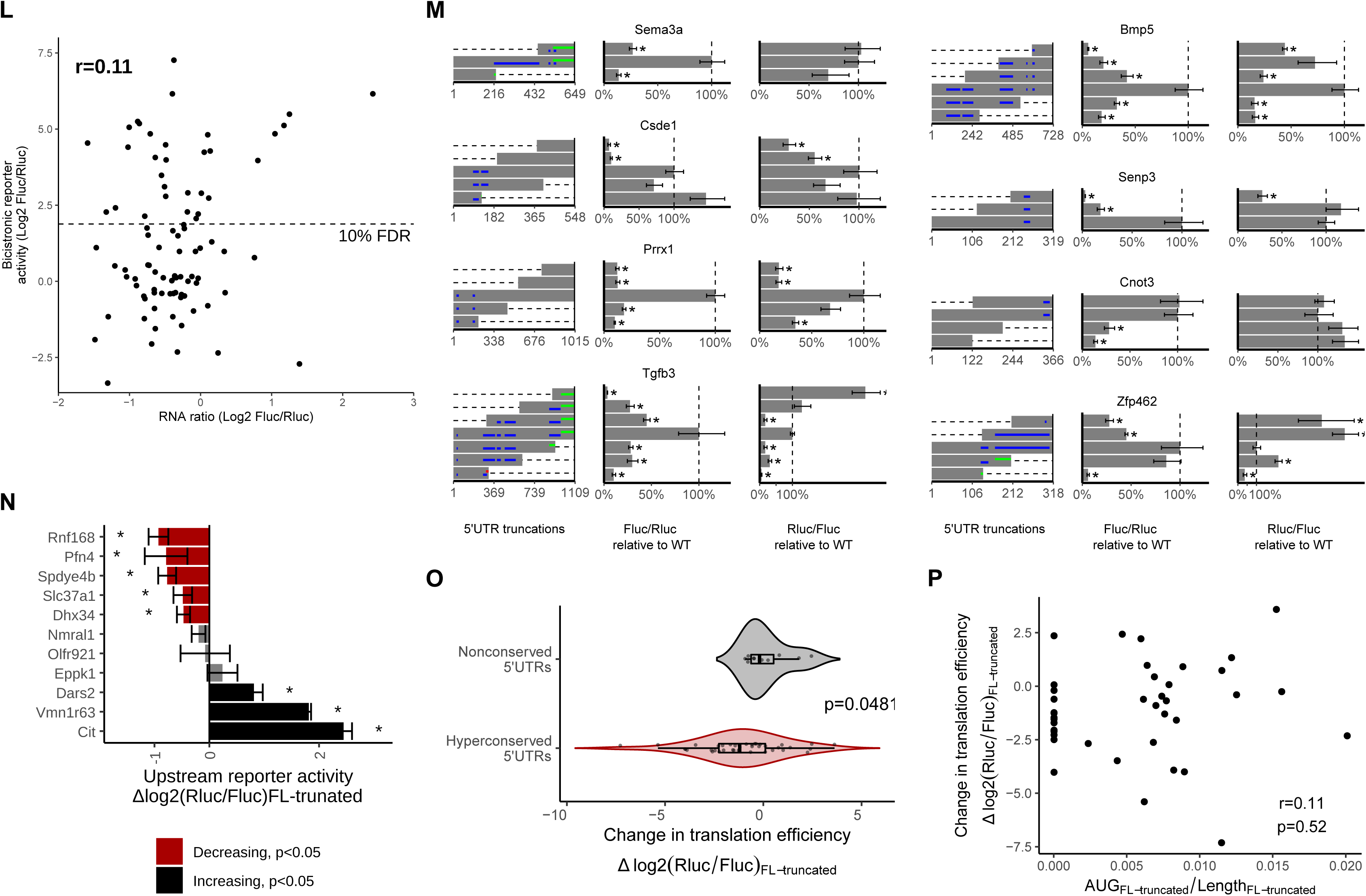
**Non-canonical translation initiation enhancer elements are a feature of hyperconserved 5’UTRs** A) Density plots of non-canonical translation initiation activities from h5UTRs by bicistronic reporter assay. X-axis is the luciferase reporter activity ratios. Jittered dots mark individual reporter ratios for each h5UTR in each cell type. B) Summarized plot of ribosome load (sum of % mRNA times the ribosome number for each fraction) differential ratio between NSCs and ESCs calculated from polysome profiles for each gene shown in **Fig. S3C-K**. Red indicates significant increase in NSCs and black indicates significant decrease (t-test p≤0.05, marked by asterisk). C) - L) Endogenous polysome profiles of NSCs versus ESCs for genes with h5UTRs that show high non-canonical translation reporter activities in NSCs compared to ESCs. Distribution of mRNAs across sucrose gradient fractions are plotted. Y-axis plots the mean percent mRNA. Error bars indicate standard error. Asterisk indicates t-test p≤0.05 for each fraction between the knockout and the wild-type. Indicated p-value (p_f_) is calculated by Fisher’s method across all fractions. Note that L) shows the profile of 18S rRNA, which indicates lower global translation in NSCs compared to ESCs. L) Scatter plot showing comparison of luciferase activity ratios versus RNA level ratios (mean from N=3) observed for the bicistronic reporters of 89 h5UTRs measured in 10T1/2 cells. Dashed line marks the 10% FDR used in Fig. 3A. M) The effect of various truncations of the h5UTRs on non-canonical initiation and total translation efficiency. These are additional data to Fig. 3D. Left: positions of truncations. Dashed lines indicate truncations and bars indicate the remaining sequences. Blue horizontal lines within bars indicate uORFs; green and red lines within bars indicate in-frame and out-of-frame uAUGs, respectively. Middle: non-canonical initiation efficiency. Right: total translation efficiency. X-axis indicates the mean luciferase reporter ratios relative to the wild-type. Error bars indicate standard error. Dashed line marks the reporter ratio for the wild-type 5’UTR. N=4. Asterisk indicates t-test p≤0.05 for each truncation mutant versus the wild-type. N) Comparison of translational activities between the full-length long, non-conserved 5’UTRs versus the only first 300nt truncation. 11 different pairs are tested. Error bars indicate standard error. Bars colored in red indicate significantly reduced translation in the shorter, truncated 300nt fragment; black indicates significant increase (t-test p≤0.05, marked by asterisk). O) Violin plot of full-length/truncated reporter activity ratios (log2) from hyperconserved and non-conserved 5’UTRs. p indicates Wilcoxon rank sum test p-value. P) Scatter plot of change in translation efficiency between full-length and truncated h5UTRs shown in Fig. 3E versus change in uAUG density (change in number of AUGs / change in length between each pair of full-length and truncated h5UTRs). r indicates pearson’s correlation coefficient and p indicates the p-value.

**Figure S4.**
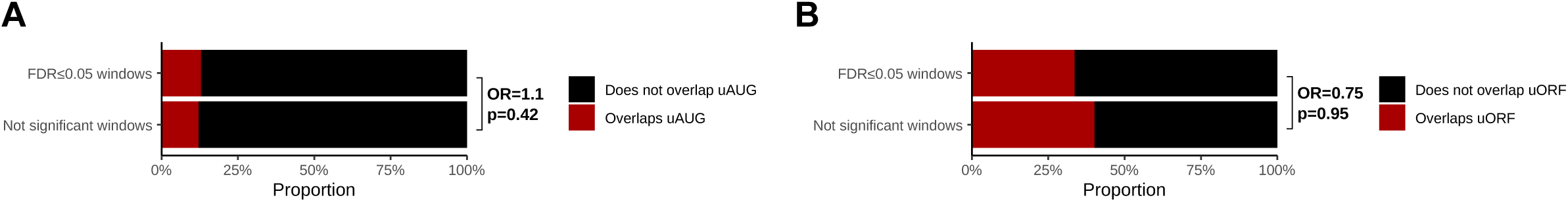
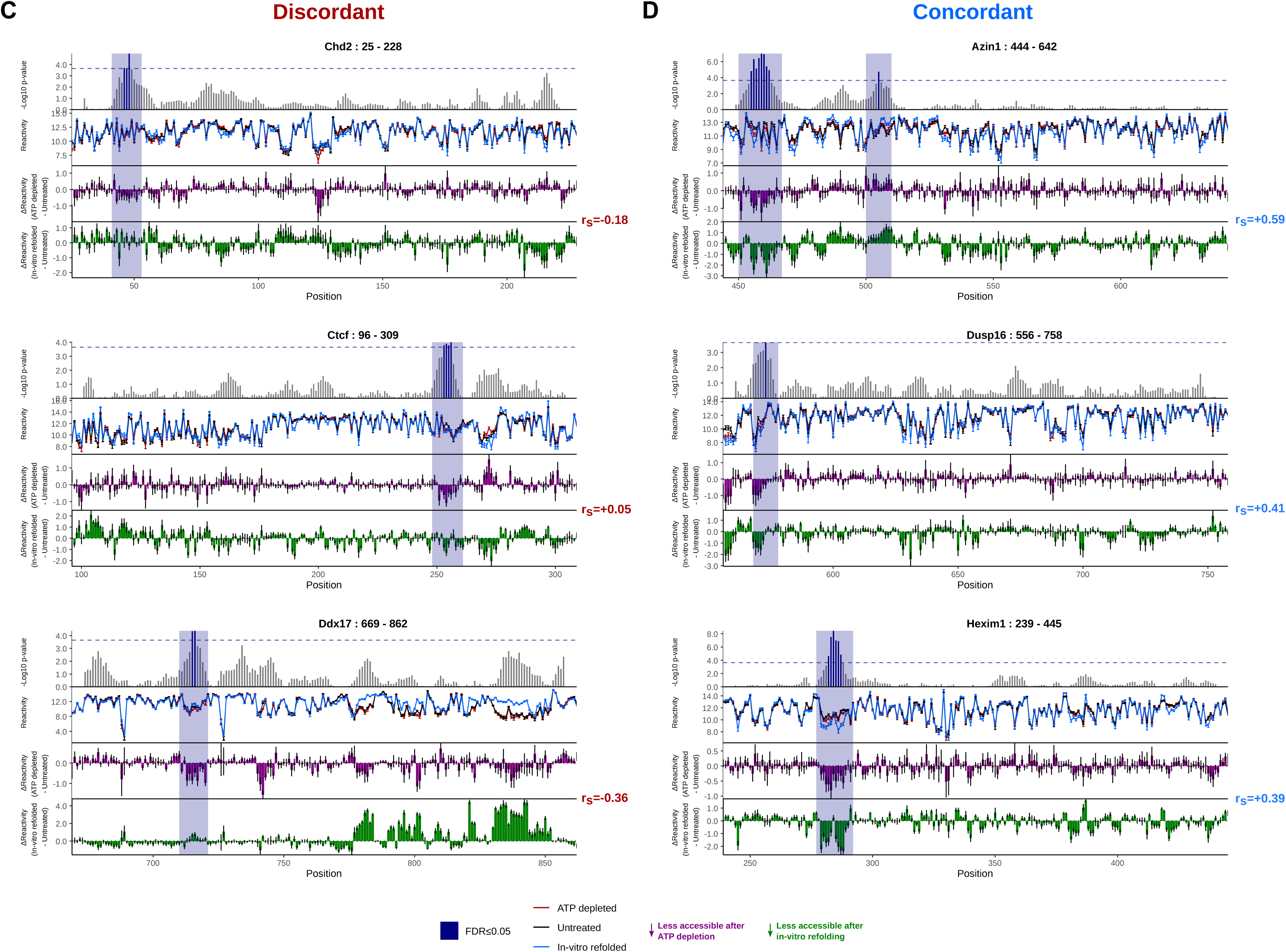
**Cellular remodeling of RNA structures in hyperconserved 5’UTRs** A) Stacked bar plots showing proportions of significant (FDR≤0.05) or not significant windows that overlap uAUG in black versus that do not overlap uAUG in red. OR indicates odds ratio for overlaps uAUG / does not overlap uAUG, and p indicates Fisher’s test p-value (one-sided, H_a_=odds ratio>0). B) Stacked bar plots showing proportions of significant (FDR≤0.05) or not significant windows that overlap uORF in black versus that do not overlap uORF in red. OR indicates odds ratio for overlaps uORF / does not overlap uORF, and p indicates Fisher’s test p-value (one-sided, H_a_=odds ratio>0). C) Zoomed-in view of differential accessibilities along h5UTRs with one or more significantly different windows under ATP depletion. Top plot shows -log10 p-value for each window. Highlighted boxes mark significantly different windows, above the dashed line indicating 5% FDR. Middle plot shows differential accessibility on the y-axis, where greater than zero indicates increased accessibility upon ATP depletion and less than zero indicates decreased accessibility. Bottom plot shows differential accessibility for in vitro refolded RNA. The three profiled regions shown here exhibit discordant profiles between accessibility changes observed in cells following ATP depletion and accessibility changes observed for in cell versus in vitro refolded RNA. D) Same as Fig. S4A, except the three profiled regions shown exhibit concordant profiles between accessibility changes observed in cells following ATP depletion and accessibility changes observed for in cell versus in vitro refolded RNA.

**Figure S5.**
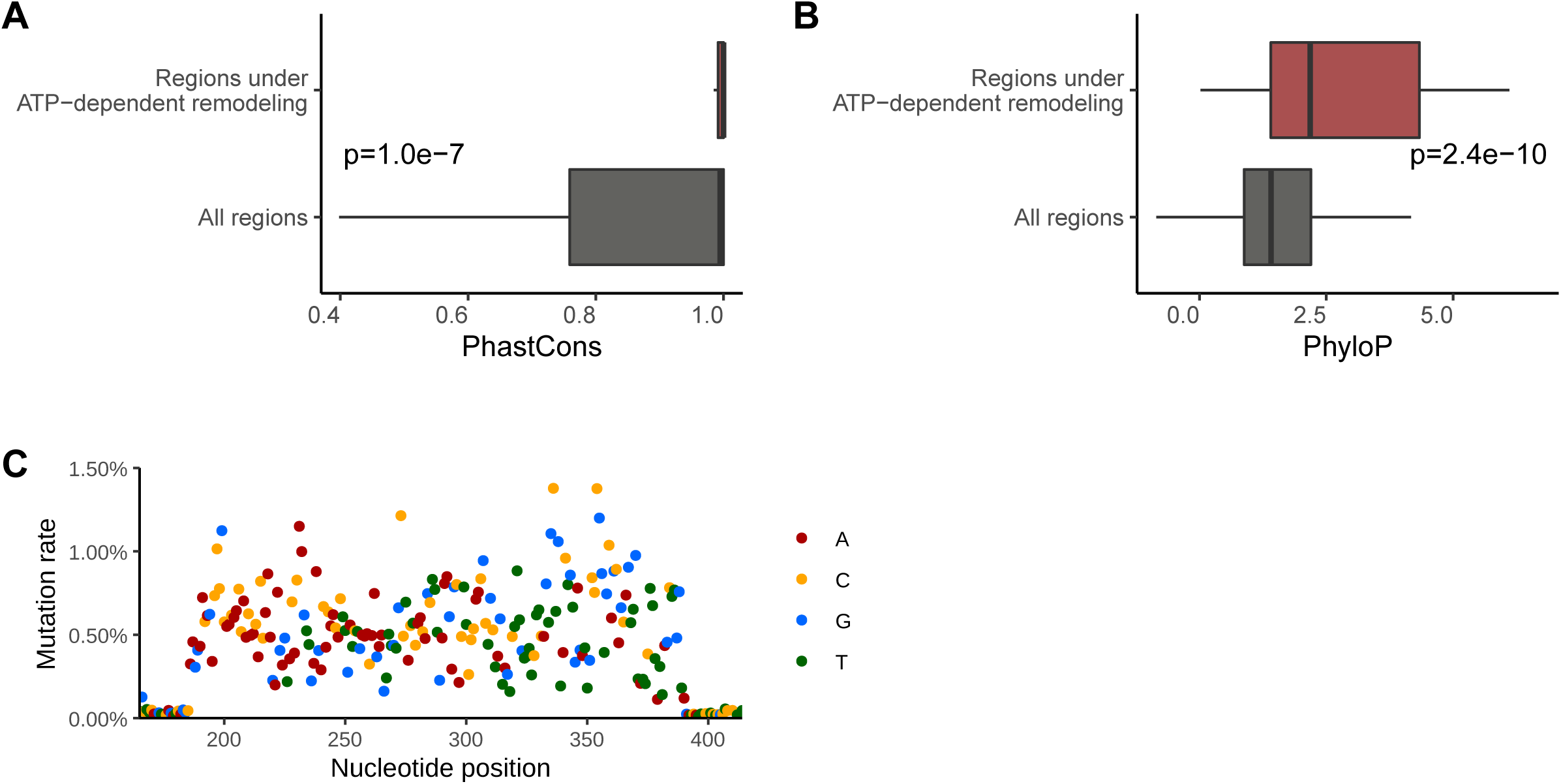
**icM2 reveals structured elements in the hyperconserved Csde1 5’UTR** A) Boxplot of average PhastCons scores in significant windows of ATP-dependent remodeling versus all windows shown in Fig. 4C. p indicates Wilcoxon rank sum test p-value. B) Same as A), but showing the distribution of average PhyloP scores. C) Plot of per-nucleotide mutation rates in the mutagenesis library used in icM^2^ analysis of Csde1 5’UTR. Dots are colored by nucleotides. Note that flanking regions have near-zero mutation rate because they are the primer regions used for amplicon sequencing. See In-cell mutate-and-map section in methods.

**Figure S6.**
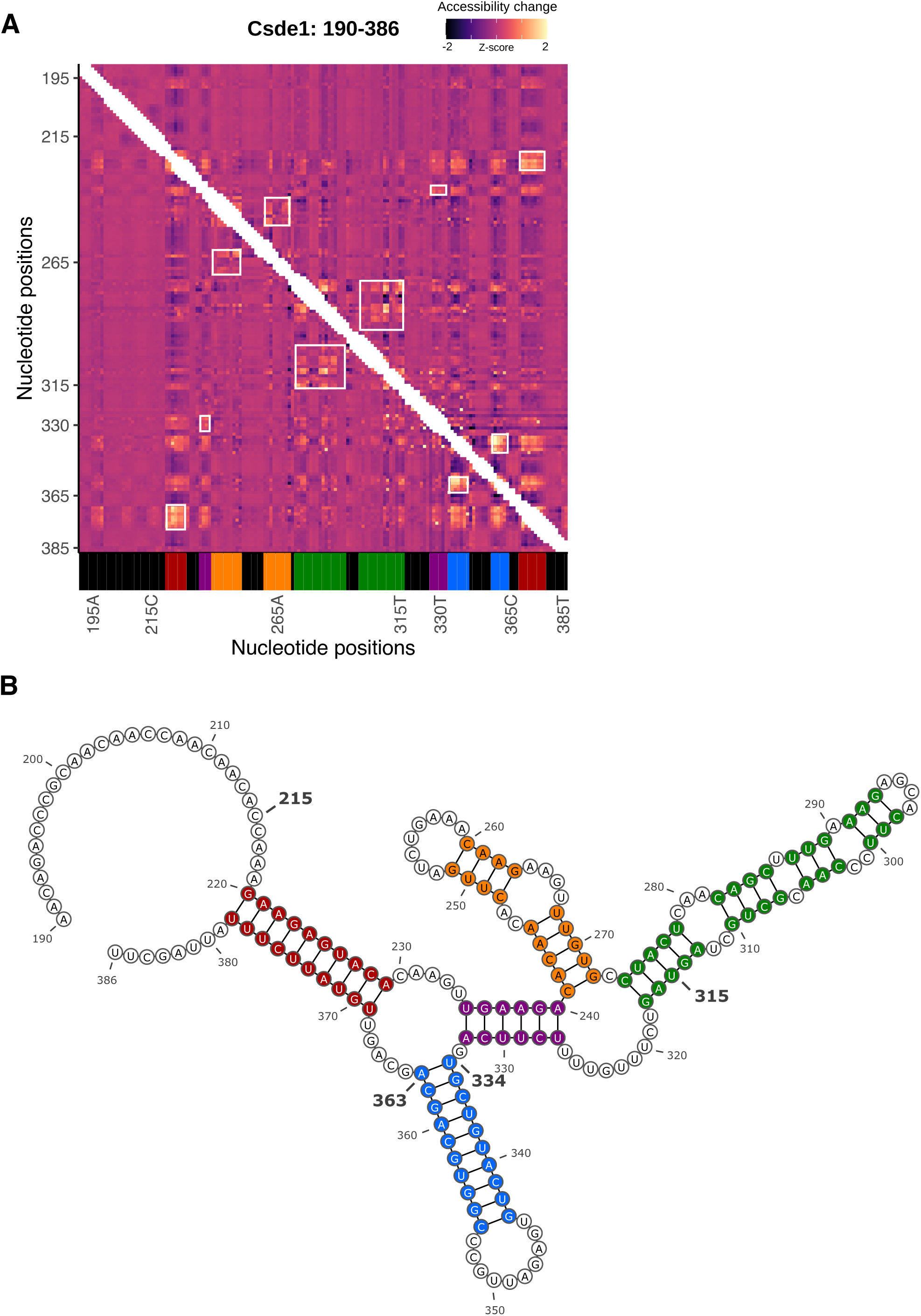
**Csde1 5’UTR tunes translation efficiency by encoding multiple alternative structures that are actively maintained by RNA helicases** A) Heatmap of in-vitro M^2^ accessibility matrix for Csde1 5’UTR from position 190 to 386. For each row, the chemical mapping profile of a single-nucleotide variant of the RNA is plotted across the columns, where the colors indicate z-scaled accessibility change values from the wild-type RNA. 1D data from each mutant are vertically stacked to display a 2D matrix. White boxes mark perturbation signals that support the model shown in **Fig. S3B**; color bars at the bottom indicate the nucleotide positions of the stems that match the same color in the model. B) The model for the in-vitro structure of Csde1 5’UTR from position 190 to 386. Also see **Fig. S3A**.

**Figure S7.**
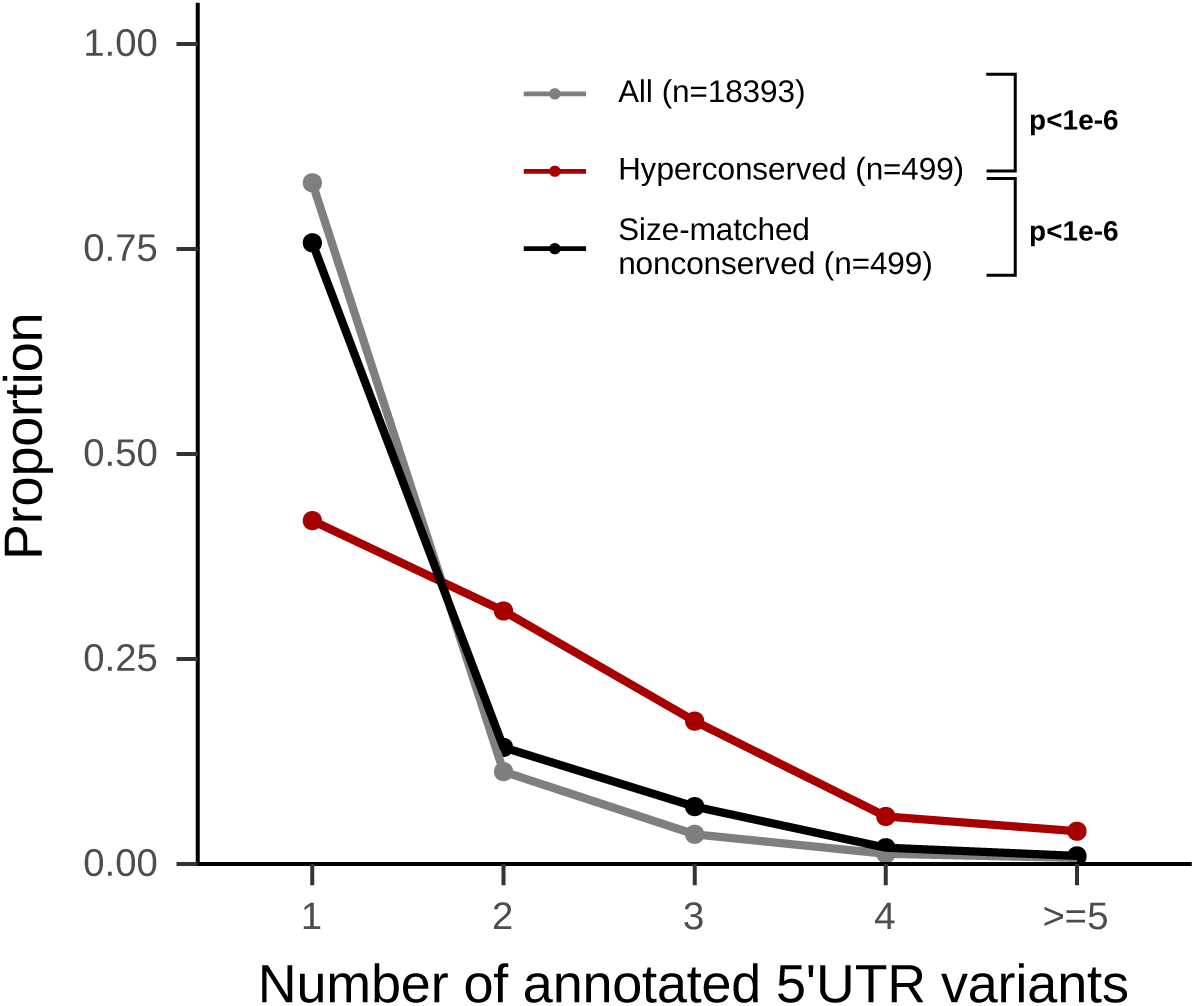
**More alternative 5’UTRs are annotated for h5UTR genes compared to non-conserved 5’UTRs** Distributions of the number of annotated alternative 5’UTRs for all genes, genes with h5UTRs, or genes with size-matched non-conserved 5’UTRs. Indicated p-values are from Wilcoxon rank sum tests for the number of alternative 5’UTRs between h5UTR genes and all genes or between h5UTR genes and size-matched non-conserved controls.

## Methods

### Data sources

The following lists the version and sources of publicly available data used in this study. RefSeq: release 84, https://ftp.ncbi.nlm.nih.gov/refseq/; 60-way PhastCons: UCSC mm10, http://hgdownload.soe.ucsc.edu/goldenPath/mm10/phastCons60way/, http://hgdownload.soe.ucsc.edu/goldenPath/mm10/database/; 60-way multiple sequence alignment: UCSC mm10, http://hgdownload.soe.ucsc.edu/goldenPath/mm10/multiz60way/; GTEX RNA-seq: V8 (GENCODE V25 annotation), https://www.gtexportal.org/home/datasets, ; GO term mapping: Bioconductor 3.10 org.Mm.eg.db, http://bioconductor.org/packages/3.10/data/annotation/html/org.Mm.eg.db.html; ENCODE: https://www.encodeproject.org/95,96,97,98,99.

### h5UTR definition

PhastCons elements represent segments of the alignment belonging to the conserved state of the phylo-HMM^27^. Each element has log odds score (LOD) = log probability under the conserved model - log probability under the non-conserved model. We downloaded the 60-way vertebrate PhastCons elements from UCSC mouse genome database, and LOD≥500 elements were subsetted. LOD≥500 selects for about 92th percentile and above for all predicted PhastCons elements, and is essentially arbitrary but chosen to pick a small enough set such that lowest scores appear very extreme upon individual inspection, while providing a meaningful enough number of 5’UTRs to allow statistical confidence. Finally, for each mouse RefSeq transcript record, the total number of 5’UTR, CDS, and 3’UTR nucleotides overlapping LOD≥500 elements are calculated (**Table S1**). 5’UTRs with ≥250nt overlap are labeled hyperconserved. As a crosscheck of the RefSeq annotation, transcription start sites of these h5UTRs are compared with ENCODE RAMPAGE datasets and found to overlap at least one TSS peak at a rate of 82%.

### GTEx data processing and analysis

Proteome-transcriptome correlations: we obtained cross-tissue proteome-transcriptome correlations from quantitative proteomics analysis of 32 human tissues that match GTEx transcriptomics samples as reported^100^. This measures how well the variation in protein levels across the tissues are predicted by variation in their RNA levels.

RNA and protein abundance/variance: we applied TMM scaling^101^ on GTEx RNA-seq count matrix. Log TMM scaled counts were quantile normalized and per-tissue median value across individuals are taken as the RNA expression level. Mean and variance of RNA expression levels are calculated from the per-tissue median values. Similarly for proteomics data, log of summed peptide abundances were quantile normalized and median values across individuals per tissue were taken. Mean and variance of protein expression levels are calculated from the per-tissue median values.

We began with a table of correlation values, RNA mean expression, RNA variance, protein mean expression and protein variance for each human gene. For a gene to be labeled hyperconserved in this table, it must have a homologous RefSeq annotated 5’UTR between humans and mice that overlap LOD≥500 PhastCons elements by at least 250nt in mouse and 150nt in humans. In other words, the hyperconserved 5’UTR must be known to both human and mouse RefSeq transcript annotations. For a gene to be labeled non-conserved in this table, no LOD≥500 PhastCons elements must overlap any 5’UTR annotation for the gene in both human and mouse transcript annotations. To be considered for analysis, each gene’s expression must be detected in at least 10 tissues for both protein and RNA. The total number of genes available for analysis is 8598. Size-matching or variance-matching non-conserved controls are selected as follows. The table is first ranked by maximum 5’UTR length annotated for the gene. For each h5UTR gene, the gene closest in 5’UTR length or RNA variance rank to the h5UTR but has a non-conserved label is selected without replacement.

### GO term enrichment analysis

Enrichment analysis is performed using R package topGO^102^. GO term-gene mappings are obtained from Bioconductor annotation package org.Mm.eg.db. Only terms with at least 10 genes annotated and at least 1 gene mapping to h5UTRs or size-matched non-conserved gene sets are tested. First, Fisher’s exact test p-value is calculated for each term, and those with Benjamini-Hochberg FDR estimate ≤0.05 are retained. topGO weight01 p-value ≤0.05, observed/expected ratio ≥=3, and minimum number of genes mapping ≥=3 is used to further filter the term list. For the final set of GO terms, semantic similarity adjacency matrix is calculated using Lin method in R package GOSemSim^103^. Adjacency matrix is ranked on the column. The rank matrix is compared with the transposed rank matrix and higher rank is taken. Network with terms as nodes are constructed and terms with maximum rank (total number of terms - 1) is connected. Clustering is performed with cluster_edge_betweenness algorithm in R package iGraph^104^.

### CRISPR knockout of hyperconserved sequences

Genome editing of E14 ESCs were achieved by using CRISPR/Cas9 nuclease-mediated recombination. sgRNAs were designed using CRISPOR^105^. sgRNA sequences were synthesized as ssDNA oligos and were cloned into the BbsI-digested expression plasmid bearing both sgRNA scaffold backbone (BB) and Cas9 nuclease, pX330-U6-Chimeric_BB-CBh-hSpCas9^106^. ∼0.5×106 cells were plated onto a single 6-well plate. After 4 hours, 1.25ug each of the plasmids carrying sgRNA pairs were transfected using 2.5uL P3000 reagent and 12uL Lipofectamine 3000 (ThermoScientific, L3000001). 12 hours after transfection, media was changed to puromycin (ThermoScientific, A1113803) containing media at 1ug/mL. After 24 hours of puromycin selection, cells were washed with PBS, trypsinized, and plated at 1000 cells/10 cm plate. 10 days later, single colonies were picked and replica plated to 2×96 well plates. One plate was used for genotyping. For 3T3 cells, the transfection and selection were performed using the same methods, but the cells were plated at limiting dilution of 0.5 cells/well into 96 well plate for expansion and split for genotyping. Cells in the genotyping plate were lysed by removing the media, adding 100uL 50mM NaOH per well, and heating at 95°C for 10min. After cooling to room temperature, 500uL 500mM Tris-HCl pH 8.0 was added to neutralize and 1:100 dilution was taken for genotyping PCR. Genotyping PCR reaction is as follows: 1x MyTaq HS Red Mix (BIO-25047), 300nM forward primer, 300nM reverse primer, 1uL 1:100 diluted crude lysate in 10uL total reaction volume. Cycling conditions are: 95°C 3min initial denaturation, followed by 30 cycles of 95°C 15s, 68°C 15s, 72°C 30s. Clones with expected shorter amplicons were further expanded. DNA from expanded clones was isolated with Wizard Genomic DNA Purification kit (Promega, A1120). Genotyping PCR reaction from expanded clones are as follows: 0.02U/uL Kapa HiFi HotStart polymerase (Roche, KR0369), 1x Kapa HiFi HotStart buffer, 300uM dNTP each, 300nM forward primer, 300nM reverse primer, 10ng gDNA in 20uL reaction. Cycling conditions are: 95°C 3min initial denaturation, followed by 30 cycles of 98°C 20s, 68°C 15s, 72°C 30s. The amplicons were Sanger sequenced at Quintara Biosciences.

### Cell culture

mESC: E14 mESCs were cultured on 0.1% gelatin-coated dishes using the following media recipe: Knockout DMEM (ThermoFisher Scientific, 10829018), 15% Embryomax FBS (MilliporeSigma, ES-009-B), 2 mM non-essential amino acids (MilliporeSigma, TMS-001-C), 2 mM L-Glutamine (MilliporeSigma, TMS-002-C), 0.1 mM 2-mercaptoethanol (ThermoFIsher Scientific, 21985023), and 10^3^ U/mL mLIF (MilliporeSigma, ESG1107; Gemini 400-495).

10T1/2: C3H10T1/2 cells were grown using the following media recipe: DMEM (ThermoFisher Scientific, 11965118), 10% FBS (ThermoFisher Scientific 26140079), 1X penicillin-streptomycin (MilliporeSigma, TMS-AB2-C).

NSC: NSC were differentiated from E14 mES cells^107^. After differentiation, NSC were grown using the following media recipe: 50% DMEM-F12 (ThermoFIsher Scientific, 11320082), 50% Neurobasal medium (ThermoFisher Scientific, 21103049), 1x modified N2 (ThermoFisher Scientific, 17502048), 1x B27 supplement (ThermoFisher Scientific, 17504044), 0.0007% BSA Fraction V (ThermoFisher Scientific, 15260037), 0.01 ug/mL murine EGF (ThermoFisher Scientific, PMG8044), 0.01 ug/mL murine FGF-basic (ThermoFisher Scientific, PMG0031), 1X penicillin-streptomycin (MilliporeSigma, TMS-AB2-C). Accutase (MilliporeSigma, A6964 was used for dissociation of NSCs.

Neuron: Neurons were differentiated from mESC derived NSCs^107^. 1-2×10^3^ NSCs were plated onto poly-ornithine/laminin coated 96-well plates using the following differentiation media recipe: 25% DMEM-F12 (ThermoFIsher Scientific, 11320082), 75% Neurobasal medium (ThermoFisher Scientific, 21103049), 1x modified N2 (ThermoFIsher Scientific, 17502048), 1x B27 supplement (ThermoFisher Scientific, 17504044), 0.005% BSA Fraction V (ThermoFisher Scientific, 15260037), 0.01 ug/mL murine FGF-basic (ThermoFisher Scientific, PMG0031). Half of the media was replaced every 2 days. After 7 days, the media was exchanged for further neuronal differentiation and maturation to: 25% DMEM-F12 (ThermoFIsher Scientific, 11320082), 75% Neurobasal medium (ThermoFisher Scientific, 21103049), 0.25x modified N2 (ThermoFIsher Scientific, 17502048), 0.25x B27 supplement (ThermoFisher Scientific, 17504044), 0.0007% BSA Fraction V (ThermoFisher Scientific, 15260037).

Limb mesenchyme culture: Stage E11.5 mouse embryos were isolated and washed in PBS. The limbs were dissected in dissection media: DMEM-F12 (ThermoFisher Scientific, 11320082), 10% FBS (ThermoFisher Scientific, 26140079), 1X penicillin-streptomycin (MilliporeSigma, TMS-AB2-C). Dissected limbs were trypsinized in 1% (w/v) trypsin (ThermoFisher Scientific, 27250018) in HBSS (ThermoFisher Scientific, 14175095) at 37°C, 5% CO_2_ incubator for 30 min. After trypsinization, tissue was resuspended in dissection media, passed through a cell strainer and 2.5X10^4^ -3×10^4^ cells per 96 well plate was cultured in dissection media overnight before transfection.

Embryoid body: E14 mESCs were trypsinized and plated on low-attachment dish at 5×10^6^ per 10cm in mESC media without LIF: Knockout DMEM (ThermoFisher Scientific, 10829018), 15% Embryomax FBS (MilliporeSigma, ES-009-B), 2 mM non-essential amino acids (MilliporeSigma, TMS-001-C), 2 mM L-Glutamine (MilliporeSigma, TMS-002-C), 0.1 mM 2-mercaptoethanol (ThermoFIsher Scientific, 21985023).

Retinoic acid treated mESC: E14 mESCs were seeded at 1×10^6^ cells per 10cm dish in mESC media without LIF + 10uM retinoic acid: Knockout DMEM (ThermoFisher Scientific, 10829018), 15% Embryomax FBS (MilliporeSigma, ES-009-B), 2 mM non-essential amino acids (MilliporeSigma, TMS-001-C), 2 mM L-Glutamine (MilliporeSigma, TMS-002-C), 0.1 mM 2-mercaptoethanol (ThermoFIsher Scientific, 21985023), 10uM retinoic acid (MilliporeSigma, R2625). Media was changed every 24 hours and cells were harvested for polysome profiling after 3 days. For the initial time course to check expression of targeted h5UTR genes, 1.5×10^5^ cells were seeded per one 6-well. Media was changed every 24 hours until lysis with Trizol (ThermoFisher Scientific, 15596026) and RNA extraction from aqueous phase on silica column (Zymo, R1013). 100ng RNA was reverse transcribed using iScript reverse transcriptase (Biorad, 1708890) in a 10uL reaction. qPCR was performed using Ssoadvanced Universal SYBR Green Supermix (Biorad, 1725270) with 2uL of 1:4 diluted reverse transcription reaction and primer pairs targeting mouse Actb and targeted h5UTRs. 2^ΔΔCq^ values relative to Actb and maximally expressed time point for each h5UTR is plotted.

### Polysome profiling

Cell culture media was replaced with cycloheximide (MilliporeSigma, C7698-1G) containing media at 100ug/mL. After 2 minutes, cells were washed, trypsinized and harvested using PBS, trypsin, and culture media containing 100ug/mL cycloheximide. ∼10×10^6^ cells were resuspended in 400uL of following lysis buffer on ice for 30min, vortexing every 10min: 25mM Tris-HCl pH 7.5, 150mM NaCl, 15mM MgCl_2_, 1mM DTT, 8% glycerol, 1% Triton X-100, 100ug/mL cycloheximide, 0.2U/uL Superase-In RNase inhibitor (ThermoFisher Scientific, AM2694), 1x Halt protease inhibitor cocktail (ThermoFisher Scientific, 78430), 0.02U/uL TURBO DNase (ThermoFisher Scientific, AM2238). After lysis, nuclei were removed by two step centrifuging, first at 1300g for 5min and second at 10000g for 5min, taking the supernatants from each. 25%-50% sucrose gradient was prepared in 13.2mL ultracentrifuge tubes (Beckman Coulter, 331372) using Biocomp Gradient Master with the following recipe: 25 or 50% sucrose (w/v), 25mM Tris-HCl pH 7.5, 150mM NaCl, 15mM MgCl_2_, 1mM DTT, 100ug/mL cycloheximide. The lysate was layered onto the sucrose gradient and ultracentrifuged on Beckman Coulter SW-41Ti rotor at 40000rpm for 150min at 4°C. The gradient was density fractionated using Brandel BR-188 into 16×750uL fractions, and 50pg in vitro transcribed luciferase RNA was added as a spike-in normalizer to each fraction. 700uL of each fraction was mixed with 100uL 10% SDS, 200uL 1.5M sodium acetate, and 900uL acid phenol-chloroform, pH 4.5 (ThermoFisher Scientific, AM9720), heated at 65°C for 5min, and centrifuged at 20000g for 15min at 4°C for phase separation. 600uL aqueous phase was mixed with 600uL 100% ethanol and RNA was purified on silica columns (Zymo, R1013). For each fraction, up to 5ug RNA was DNase treated at 37°C for 30min using 0.2U/uL TURBO DNase with 1U/uL Superase-In in 30uL and purified again on silica column. 100ng RNA from each fraction was reverse transcribed using iScript reverse transcriptase (Biorad, 1708890) in a 10uL reaction. qPCR was performed using Ssoadvanced Universal SYBR Green Supermix (Biorad, 1725270) with 2uL of 1:4 diluted reverse transcription reaction and primer pairs targeting the HCE knockout h5UTR genes or the spike-in luciferase. Ct values of target genes were normalized to Ct values of spike-in luciferase and plotted as proportions across the 16 fractions. T-statistic and p-value is calculated for difference in means between the two genotypes from N=4∼6 for each fraction. Fisher combined p-value is calculated for no difference across all fractions.

### Reporter constructs for luciferase assays

Bicistronic reporter gateway plasmid, pRF_gwy, is constructed from pRF vector which has SV40 promoter and two reporter genes, Renilla luciferase and Firefly luciferase, with multiple cloning sites in between them^108^. Gateway cassette A (ThermoFisher Scientific, 11828029) was inserted in between Renilla and Firefly luciferases, replacing the cloning sites using two EcoRI sites.

RNA normalizing reporter gateway plasmid, pRF_D1, is constructed from pRF vector by replacing the cloning sites with HCV IRES and inserting gateway cassette A in between AvrII and EcoRV sites upstream of Renilla luciferase. Downstream of the Renilla luciferase is the Firefly luciferase and the HCV IRES between them such that the HCV IRES-translated downstream Firefly luciferase normalizes for differences in RNA levels to enable measurement of translation efficiency.

Full-length h5UTRs were synthesized and cloned into pENTR1A (ThermoFisher Scientific, A10462) by SGI. Truncation variants are cloned by PCR from synthesized full-length sequences into pENTR1A vectors. Gateway LR Clonase II (ThermoFisher Scientific, 11791020) is used to recombine the full-length or truncation variants into either pRF_gwy or pRF_D1 vectors. Mutant Csde1 5’UTRs were cloned by Gibson assembly reaction (NEB, E2621S) using mutation containing ssDNA templates with homology arms and two upstream/downstream fragments.

Full-length wild-type and mutant Csde1 5’UTRs were inserted pGL3 (Promega, E1751) plasmid in between EcoRI and NcoI sites upstream of Firefly luciferase gene. In vitro transcription template was amplified with T7 promoter sequence containing primer and in vitro transcribed using T7 RNA polymerase (NEB, E2040S). The IVT RNAs were capped using Vaccinia virus capping enzyme (Cellscript, C-SCCE0625) and polyA tailed using polyA polymerase (Cellscript, C-PAP5104H).

### Reporter transfection for luciferase assays

For DNA transfections, 200ng plasmid DNA is transfected to cells plated on 96-well plates. For 10T1/2 cells, mESCs, NSCs, limb mesenchyme culture, and embryoid bodies, 0.5uL Lipofectamine 2000 (ThermoFisher Scientific, 11668030) is used per one well. For neurons, 0.2uL Viafect (Promega, E4981) is used per well. Cells were incubated for 4 hrs with transfection reagent, DNA in OptiMEM media (ThermoFisher Scientific, 31985062). Cells were then washed with PBS, and the media was changed back to regular growth media.

For RNA transfection, 200ng Firefly luciferase RNA and 10ng Renilla luciferase RNA is transfected to cells plated on 96-well plates. 0.5uL Lipofectamine 2000 is used per one well.

### Luciferase assays

For DNA transfections, cells were lysed using Passive Lysis Buffer (Promega, E1941) for 30min at room temperature, 48 hours after transfection. For RNA transfections, cells were lysed 6 hours after transfection. Firefly and Renilla luciferase values were read using Dual-Glo Luciferase Assay System (Promega, E2920) for >96 samples or Dual-Luciferase Reporter Assay System (Promega, E1910) for fewer samples, on Promega GloMax-Multi plate reader. In all experiments, log ratios of the two luciferase activities are taken for each well.

For the large-scale bicistronic reporter assays across multiple cell types, the data are quantile normalized across all replicate samples. normalmixEM function from R package mixtools is used for mixture modeling of maximum replicate-average values^109^. False discovery estimate at a cutoff is calculated as the proportion of the mixture distribution above the chosen cutoff that comes from the lower component. For identification of h5UTRs with significant differential activity across cell types, F-statistic is calculated, and Benjamini-Hochberg procedure is used with their p-values to estimate the FDR. N=4 for C10T1/2, mESC and EB; N=6 for NSCs, neurons, limb mesenchyme culture.

For other reporter assays, all statistics are calculated from log ratios of the luciferase activities. Mean and error bars when plotted in linear scale are back transformed from the mean and standard errors of the log scale values.

### In-cell DMS probing following ATP depletion and multiplexed mutational profiling

100 h5UTRs were chosen for amplicon sequencing based on coverage profiles from ENCODE E14 mESC RNA-seq data. A total of 384 primer pairs were designed across these h5UTRs using ThermoAlign and following parameters (others are left at default): primer_size_range=16-40, GC_range=25-75, Tm_range=64-72, primer_conc=300, Na=0, K=75, Tris=10, Mg=2, dNTPs=1.2, amplicon_size_min=250, amplicon_size_max=250^110^. Primers were split into 4 pools of 96 pairs using PrimerPooler^111^. A final set of 380 pairs were synthesized by Eurofins Genomics.

10×10^6^ mESCs were incubated for 10min in ATP depletion media: DMEM without glucose (ThermoFisher Scientific, A1443001), 10mM 2-Deoxy-D-glucose (MilliporeSigma, 25972), 10mM sodium azide (MilliporeSigma, 71289). The cells were washed, trypsinized and harvested using PBS, trypsin, and finally resuspended in 3.5mL mESC media all containing 10mM 2DG and 10mM NaN_3_. 1mL 1M bicine (Millipore Sigma, B3876), titrated to pH 8.5 at 25°C, is added to resuspended cells (250mM final bicine concentration). 500uL 16% dimethyl sulfate (MilliporeSigma, D186309) in ethanol is added (1.6% final concentration). Cells are mixed and incubated for 6min at 37°C. 2.5mL ice-cold 30% BME (MilliporeSigma, M3148) in ethanol is added to quench the reaction. This DMS modification protocol is adapted from protocol and data reported in^70^. Following centrifuge to remove the supernatant, the cells are lysed in Trizol (ThermoFisher Scientific, 15596026). For the untreated condition without ATP depletion treatment, all procedures are the same except for initial 10min incubation in ATP depletion media and inclusion of 2DG and NaN_3_ in all media. Three independent samples were collected for each condition. Total RNA is phase extracted with chloroform and aqueous phase is purified on silica columns (Zymo, R1013). 10-20ug RNA is DNase treated at 37°C for 30min using 0.2U/uL TURBO DNase (ThermoFisher Scientific, AM2238) with 1U/uL Superase-In (ThermoFisher Scientific, AM2696) in 60uL and purified again on silica column.

1ug RNA is mixed with 1uL 1uM 96x primer pool (96x1uM amplicon reverse primers for total of 1uM oligos) and denatured in 6.25uL total volume (with H2O) at 65°C for 2min, then chilled to 4°C. Reverse transcription reaction conditions are as follows: 20mM Tris-HCl pH7.5, 75mM KCl, 10mM MgCl2, 5mM DTT, 500nM TGIRT (InGex, TGIRT50), 1U/uL Superase-In; 19uL total reaction volume. The RT reaction is preincubated for 25°C for 30min, then initiated with addition of 1uL 12.5mM dNTP each. After incubation at 60°C for 1hour, 1uL 2.5M NaOH is added and the reaction is heated at 95°C for 3min. 1uL 2.5M HCl and 1uL 500mM Tris-HCl pH 7.5 is added to neutralize. 29uL SPRIselect beads (Beckman Coulter, B23318) is used for purification of cDNA; elution volume is 6uL. 4×96 pooled reactions for a total of 384 targets are performed for each sample. To each set of 96 pool cDNA, multiplex PCR is performed with 5uL cDNA, 0.2mM dNTP, 2uM 96x forward primer pool (96x2uM each primer for total of 2uM oligos), 2uM reverse primer pool, 1x SYBR Green I (ThermoFisher Scientific, S7563), Q5 Hot Start DNA polymerase (NEB, M0493S), 1x Q5 Hot Start Reaction Buffer, 1x Q5 Hot Start High GC Enhancer, in total 30uL reaction volume. Cycling conditions are: 98°C 30s initial denaturation, followed by 15-25 cycles (terminated before plateau) of 98°C 10s, 56°C 40s, 76°C 10s. PCR is run for 15-25 cycles. Each 96 pool multiplex PCR reaction is then used in a master mix of second PCR with 96 individual primer pair reactions: 0.2mM dNTP, 500nM forward primer, 500nM reverse primer, 1x SYBR Green I, Q5 Hot Start DNA polymerase, 1x Q5 Hot Start Reaction Buffer, 1x Q5 Hot Start High GC Enhancer, in total 6uL reaction volume. Cycling conditions are: 98°C 30s initial denaturation, followed by 20 cycles of 98°C 10s, 62°C 10s, 76°C 10s. 384 individual reactions are pooled and purified on silica columns (NEB, T1030S). Amplicon pool is end-prepared, Illumina adaptor sequences are ligated, adaptor-ligated DNA is size selected with SPRIselect beads for 370bp, and 3 cycle barcoding PCR is performed (NEB, E7645S). 2×150bp paired end sequencing data is generated on Illumina HiSeq 4000 at Novogene.

### 1D accessibility data analysis

After bcl conversion and demultiplexing with Illumina bcl2fastq, adaptor sequences are further trimmed using cutadapt^112^. Overlapping paired-end reads are merged using BBMerge^113^. Trimmed and merged reads are aligned to indexed reference of amplicon sequences using Bowtie2 with the following parameters: -D 20 -R 3 -N 0 -L 20 -i S,1,0.50 --mp 3,1 --rdg 5,1 --dpad 30^114^. Alignments are filtered to have at least 4 mutations with minimum PHRED estimate of 30 (quality cutoff for substitutions only) and minimum length 150nt. Co-occurrence of modifications at more than 4 positions is considered highly unlikely and assumed to be PCR jackpots. Duplicates are grouped using Hamming distance and directional network algorithm for clustering as implemented for UMIs in umi_tools^115^. The consensus reads are then parsed and counted for different classes of mutations using functions in ShapeMapper 2 library^116^. This mutational profiling data processing pipeline results in a matrix of mutation counts, where rows are each nucleotide position of the amplicons and columns are the different samples.

For each amplicon, the counts are first normalized by TMM^101^. As a quality filter, we require that the average pairwise Pearson correlations between replicate samples for both conditions must be greater than 0.65 for the amplicon to be further analyzed. For statistical significance of differential mutation rates between ATP depletion and untreated conditions at each nucleotide, we use voom-limma which models mean-variance bias and calculates moderated T-statistics^117, 118^. To analyze per-window accessibility pattern differences, for each sliding window of 11nt, we calculate Anderson-Darling statistic between per-nucleotide T-statistic values of the window versus per-nucleotide T-statistic values of the whole amplicon. False discovery rates are estimated by Benjamini-Hochberg procedure. Overlapping significant windows above the chosen cutoff are merged.

### In-cell mutate-and-map

Error-prone PCR was performed as follows: 0.05U/uL Mutazyme II (Agilent, 200550), 1x Mutazyme II buffer, 1x SYBR, 200uM dNTP each, 300nM forward primer, 300nM reverse primer, 100pg Csde1 5’UTR fragment; 95°C 2min initial denaturation, 10 cycles of 95°C 30s, 63°C 20s, 72°C 1min. 100pg input amount was determined by initially varying the input amounts to determine the amount at which the PCR is in exponential phase (50% of signal plateau). These parameters result in a mutation rate of approximately 1 per 200nt (**Fig. S5C**). The error-prone PCR amplicon has homology arms for Gibson assembly into pcDNA5/FRT (ThermoFisher Scientific, V601020) plasmid with EGFP. The final construct has a flanking primer region for cDNA amplification upstream of the mutagenized 5’UTR and EGFP open reading frame downstream. NEB 10-beta *E. coli* cells (NEB, C3020K) are transformed by electroporation and plated over a total of 2000cm2 area to give ∼10000 colonies. The plate is scraped and the mutagenesis library plasmid pool is purified.

mESCs are grown, dissociated, and resuspended at 5×10^5^ cells/mL. 3ug mutagenesis library plasmid pool DNA and 7.5uL Lipofectamine 2000 in 100uL OptiMEM (ThermoFisher Scientific, 31985062) are mixed with 2mL cells (1×10^6^ cells) in mESC media. The cells are incubated for 10min in suspension, centrifuged, washed with mESC media, and plated into one well in a 6-well plate. 24 hours after transfection, the cells are washed, trypsinized and resuspended in 3.5mL mESC media. 1mL 1M bicine (Millipore Sigma, B3876), titrated to pH 8.5 at 25°C, is added to resuspended cells (250mM final bicine concentration). 500uL 16% dimethyl sulfate (MilliporeSigma, D186309) in ethanol is added (1.6% final concentration). Cells are mixed and incubated for 6min at 37°C. 2.5mL ice-cold 30% BME (MilliporeSigma, M3148) in ethanol is added to quench the reaction. Following centrifuge to remove the supernatant, the cells are lysed in Trizol (ThermoFisher Scientific, 15596026). Total RNA is phase extracted with chloroform and aqueous phase is purified on silica columns (Zymo, R1013). 10-20ug RNA is DNase treated at 37°C for 30min using 0.2U/uL TURBO DNase (ThermoFisher Scientific, AM2238) with 1U/uL Superase-In (ThermoFisher Scientific, AM2696) in 60uL and purified again on silica column.

1ug RNA is mixed with 1uL 1uM RT primer targeting the downstream EGFP and denatured in 6.25uL total volume (with H2O) at 65°C for 2min, then chilled to 4°C. Reverse transcription reaction conditions are as follows: 20mM Tris-HCl pH7.5, 75mM KCl, 10mM MgCl2, 5mM DTT, 500nM TGIRT (InGex, TGIRT50), 1U/uL Superase-In, 19uL total reaction volume. The RT reaction is preincubated for 25°C for 30min, then initiated with addition of 1uL 12.5mM dNTP each. After incubation at 60°C for 1hour, 1uL 2.5M NaOH is added and the reaction is heated at 95°C for 3min. 1uL 2.5M HCl and 1uL 500mM Tris-HCl pH 7.5 is added to neutralize. 29uL SPRIselect beads (Beckman Coulter, B23318) is used for purification of cDNA; elution volume is 7uL. For each of the 3 tiling primer pairs, PCR reaction is performed as follows: 0.2mM dNTP, 300nM forward primer, 300nM reverse primer, 1x SYBR Green I (ThermoFisher Scientific, S7563), Q5 Hot Start DNA polymerase (NEB, M0493S), 1x Q5 Hot Start Reaction Buffer, 1x Q5 Hot Start High GC Enhancer, 2uL cDNA, in total 20uL reaction volume. Cycling conditions are: 98°C 30s initial denaturation, 20 cycles of 98°C 10s, 64°C 10s, 72°C 30s. The amplicons are purified on silica columns and pooled. The pooled DNA is end-prepared, Illumina adaptor sequences are ligated, adaptor-ligated DNA is size selected with SPRIselect beads for 370bp, and 3 cycle barcoding PCR is performed (NEB, E7645S). 2×150bp paired end sequencing data is generated on Illumina HiSeq 4000 at Novogene.

### 2D accessibility data analysis and structure models

After bcl conversion and demultiplexing with Illumina bcl2fastq, adaptor sequences are further trimmed using cutadapt^112^. Overlapping paired-end reads are merged using BBMerge^113^. Trimmed and merged reads are aligned to indexed reference of amplicon sequences using Bowtie2 with the following parameters: -D 20 -R 3 -N 0 -L 20 -i S,1,0.50 --mp 3,1 --rdg 5,1 --dpad 30^114^. Alignments are filtered to have at least 4 mutations with minimum PHRED estimate of 30 (quality cutoff for substitutions only) and minimum length 150nt. The consensus reads are then parsed for different classes of mutations using functions in ShapeMapper 2 library^116^. Only substitution mutations are taken and pairwise mutation co-occurrence counts are recorded into a symmetric 2D matrix for each mutation spectrum, where rows and columns are positions of the nucleotide along the amplicon. For a detailed explanation of the mechanisms and analysis of M2 signal generated by mutational profiling, see M2-seq paper^119^.

For each amplicon, 2D matrices for each type of substitution mutations are summed. TMM factors are used to normalize the matrix per column and logarithm of normalized values are used^101^. Based on mean-variance (per column) plot, minimum count threshold is selected. We chose to mask the nucleotide positions which average count less than the threshold. This results in filtering of nearly all G nucleotides as expected, which have lower signal (lower modification-mutation rates) than other bases and different variance distribution^70^. The diagonal of the matrix within 4nt is also masked due to extremely low co-variation. The matrix is further z-scaled per row and centered by the median of the entire matrix. For visualization, the z-score values are winsorized to 2 standard deviations. For clustering similar mutant (row) accessibility patterns, missing values in the diagonals are first imputed by k-nearest neighbor method with k=10. Classical multidimensional scaling is used to translate pairwise distances (Euclidean, i.e. PCA) between the mutant accessibilities into N dimensions. The resulting points from N=2 are heuristically grouped by inspection of scatter plot visualization. Two clusters are identified and per-column average z-scores are calculated.

To model structures for region B, we subsetted positions 215-315. Cluster average scores are used as constraints for partition function calculation in Vienna RNA^120^. 250 structures are sampled for each cluster constrained partition function and used as input suboptimals to REEFFIT^86^. As the input accessibility data to REEFFIT, we processed the TMM normalized log count matrix as follows. First, all cell values within each nucleotide were scaled and centered. The diagonal of the matrix within 4nt was masked; missing values are imputed by k-nearest neighbor method with k=10. The distribution of all values in the matrix are bimodal, similar to distributions observed for log data for existing M^2^ datasets generated with SHAPE and capillary electrophoresis that are available in RMDB^121^. Since REEFFIT estimates paired and unpaired reactivity distributions by mixture modelling of exponential distributions, we rescaled the DMS icM^2^ matrix on exponential distribution such that the quantile/rank is preserved but the distribution of values closely approximate SHAPE-CE M^2^ datasets downloaded from RMDB. For visualization of the landscape, we used pairwise distance metrics, structure clustering, and medoid assignment produced by REEFFIT. Sum of weights for structures belonging to each of the 3 clusters represented by a medoid structure is presented in the main figure. We used bootstrapping to robustly estimate population fraction errors. Each bootstrap runs (200 times total) using the same parameters, except that the columns (positions) of the data matrix are shuffled. This gives bounds on the frequency of the structures in the population. Reported values in the manuscript are estimated mean±standard deviation of the weight.

## Data availability

All high-throughput sequencing data generated (related to Figures 4, S4, 5, S5, 6 and S6) are deposited to GEO with accession code GSE155656. Sources for publicly available data are described in methods. Other data that support this study’s findings are available from the authors upon reasonable request.

## Code availability

All softwares used to analyze the study data are listed in the methods section and in the Nature Research Reporting Summary and are publicly available. All codes used to analyze icM^2^ data are available through a Github repository: github.com/gwbyeon/icm2p. Additional assistance with our data processing pipeline scripts or analysis codes will be provided upon request.

